# FAM72A antagonizes UNG2 to promote mutagenic uracil repair during antibody maturation

**DOI:** 10.1101/2020.12.23.423975

**Authors:** Yuqing Feng, Conglei Li, Jessica Stewart, Philip Barbulescu, Noé Seija Desivo, Alejandro Álvarez-Quilón, Rossanna C. Pezo, Madusha L.W. Perera, Katherine Chan, Amy Hin Yan Tong, Rukshana Mohamad-Ramshan, Maribel Berru, Diana Nakib, Gavin Li, Gholam Ali Kardar, James Carlyle, Jason Moffat, Daniel Durocher, Javier M. Di Noia, Ashok S. Bhagwat, Alberto Martin

**Author notes:** Shared first author (Yuqing Feng, Conglei Li).

## Abstract

Activation-induced cytidine deaminase (AID) catalyzes the deamination of deoxycytidines within *Immunoglobulin* (*Ig*) genes to induce somatic hypermutation (SHM) and class switch recombination (CSR) ^1,2^. AID-induced deoxyuracils within *Ig* loci are recognized and processed by subverted base excision and mismatch repair pathways that ensure a mutagenic outcome in B lymphocytes ^3–8^. However, it is unclear why DNA repair pathways that remove deoxyuracil from DNA are not efficient at faithfully repairing AID-induced lesions. Here, we identified through a genome-wide CRISPR screen that FAM72A, a protein with no ascribed function, is a major determinant for the error-prone processing of deoxyuracil. *Fam72a*-deficient CH12F3-2 B cells and primary B cells from *Fam72a*^−/−^ mice have reduced CSR and SHM frequencies. The SHM spectrum in B cells from *Fam72a*^−/−^ mice is opposite to that observed in *Ung2*^−/−^ mice ^9^, suggesting that UNG2 is hyperactive in FAM72A-deficient cells. Indeed, FAM72A binds to UNG2 resulting in reduced UNG2 activity, and significantly reduced protein levels in the G1 phase, coinciding with peak AID activity. This effect leads to increased genome-wide deoxyuracils in B cells. By antagonizing UNG2, FAM72A therefore increases U•G mispairs that engage mutagenic mismatch repair promoting error-prone processing of AID-induced deoxyuracils. This work shows that FAM72A bridges base-excision repair and mismatch repair to modulate antibody maturation.

---

AID catalyzes the production of deoxyuracil (dU) through cytosine deamination at Ig loci precipitating antibody diversification processes SHM, CSR, and gene conversion in B lymphocytes ^2,10^. UNG2 normally functions in base excision repair (BER) to replace dU with dC preventing mutations. Paradoxically, UNG2 promotes transversion mutations at G:C basepairs of AID-induced dUs during SHM ^9^. The mismatch repair (MMR) pathway is also coopted during antibody diversification by producing mutations at A:T basepairs during the repair of the AID-induced U•G mispairs. The mutagenic properties of MMR during SHM seems to be caused by the disruption of canonical MMR by UNG2 ^11–13^. As the error-free repair of U•G mispairs occurs routinely in mammalian cells, it is unknown why B cells do not faithfully repair all AID-induced dUs. In addition, it is unknown how the DNA repair activity of UNG2 is subverted during SHM and CSR. These observations suggest the existence of a factor(s) within B lymphocytes that can subvert DNA repair pathways that normally repair dUs.

To uncover novel factors involved in CSR, we conducted a genome-wide pooled CRISPR screen in a murine B cell line. For this analysis, we used CH12F3-2 cells (hereafter referred to as CH12 cells), which switch from IgM to IgA after anti-CD40, IL-4 and TGF-β (CIT) cocktail stimulation ^14^. Cas9-expressing CH12 clones (**Extended data Fig. 1a**)were combined and transduced with the mouse TKO (mTKO) pooled lentiviral library consisting of 95,528 Cas9 guide (g)RNA sequences targeting 19,463 genes ^15^. Transduced cells were selected using resistance to puromycin, and then treated with CIT to stimulate switching from IgM to IgA. IgA^+ve^ and IgA^−ve^ cells were sorted by flow cytometry, and gRNA representation was compared to unsorted cells by sequencing genomic DNA from each of these populations (**Fig. 1a**). The relative paucity of gRNAs from IgA^+ve^ to unsorted populations identified a number of known genes with well-established functions in CSR, including *Aicda* (which encodes AID), *Ung,* members of the mismatch repair pathway, as well as the *Tgf-β receptor* (**Fig. 1b**). Among factors that were identified by this screen, we focused on the *Fam72a* gene (**Fig. 1b**), which is largely uncharacterized but interacts with UNG2 ^16^, although no functional consequence was established for this interaction, thus deserving further characterization.

**Figure 1.**
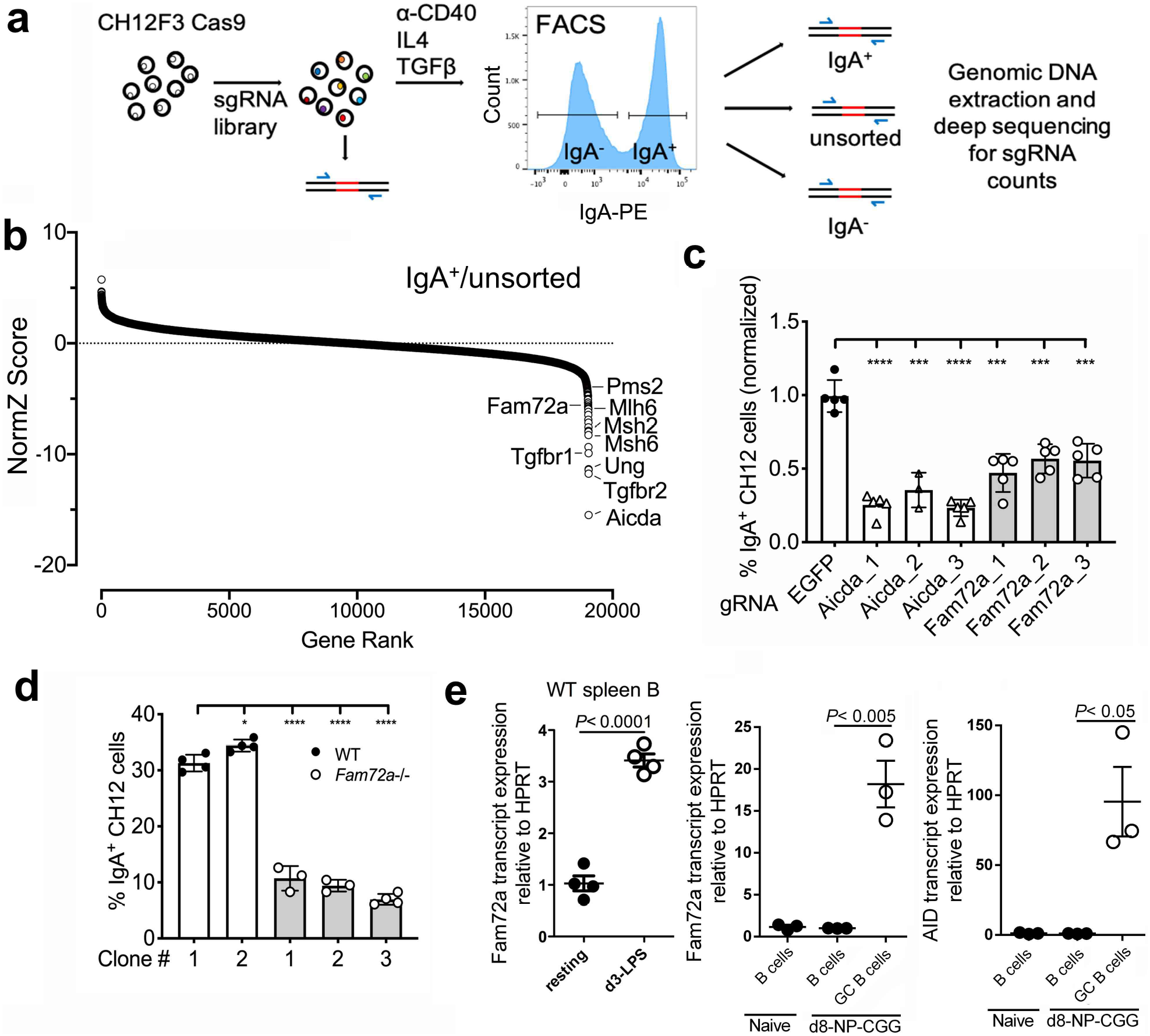
Whole-genome CRISPR screen identified FAM72A as a novel factor required for CSR. (**a**) Schematic of the CRISPR/Cas9 sorting screen in CH12 cells. CH12 cells stably expressing Cas9 was transduced with a mouse gRNA library that on average contains 5 gRNA per gene. Transduced populations were treated with CIT cocktail (CD40 ligand, IL4, TGFβ) to induce CSR from IgM to IgA, followed by staining for surface IgA and FACS sorting to separate the switched (IgA^+^) versus the unswitched (IgA^−^) population. Genomic DNA from the initially transduced cells (T0), and from unsorted, IgA^−^, IgA^+^ population was isolated, and sequenced. (**b**) gRNA was ranked using a NormZ plot (standard deviations from the mean) comparing IgA^+^ cells to the unsorted population. (**c**) Validating the role of FAM72A in CSR in Cas9-CH12 cells. A lentiviral vector carrying gRNA targeting different areas of *Fam72a* and *Aicda* genes were used to transduce CH12 cells stably expressing Cas9, followed by puromycin selection for positively transduced cells. Cells were harvested and analyzed by flow cytometry for IgA expression after 2 days post CIT treatment. Data was compared to CH12 cells expressing gRNAs targeting EGFP, which was set as 1. (**d**) Defective CSR was observed in three different *Fam72a*^−/−^ CH12 clones generated using CRISPR/Cas9 gene editing. Cells were analyzed 2 days post CIT treatment. (**e**) Murine splenic B cells were stimulated with lipopolysaccharide (LPS) *ex vivo* for 3 days, followed by qPCR for *Fam72a* and *HPRT* as control (left panel; n=4 mice per group). WT mice were immunized with NP29-CGG and Alum adjuvant and at d8 post-immunization, naïve splenic B cells (B220^+^/GL-7^−^/Fas^−^) and germinal center B cells (GC B; B220^+^/GL-7^+^/Fas^+^) were sorted by flow cytometry (with naïve splenic B cells from unimmunized WT mice as control), followed by *Fam72a* and *AID* expression assessment by qPCR (right panel; n=3 mice per group). *, p<0.05; **, p<0.01; ***, p<0.001; ****, p<0.0001.

To validate the role of FAM72A in CSR, we employed three different gRNAs to knock out *Fam72a* in CH12 cells in bulk **(Extended data Fig. 1b)**, and found that all three gRNAs efficiently reduced IgA CSR (**Fig. 1c**). We further generated *Fam72a* knockout CH12 clones (**Extended data Fig. 1c, 2a**), and found that three independent *Fam72*^−/−^ CH12 clones exhibited markedly reduced IgA CSR compared to two *Fam72a*^+/+^ wildtype (WT) CH12 clones (**Fig. 1d**). Strikingly, *Fam72a* transcription is increased ~3-fold in LPS-stimulated primary mouse B cells *ex vivo*, and ~20-fold in germinal center B cells *in vivo* (**Fig. 1e**), suggesting an important role at this stage of B cell development.

To further validate the role of FAM72A in antibody diversification, we generated *Fam72a* knockout mice (**Extended data Fig. 3**). We found that B cell profiles in bone marrow and spleens were comparable between *Fam72a*^−/−^ mice and littermate control *Fam72*^+/+^ mice (**Extended data Fig. 4, 5**), indicating that FAM72A is dispensable for B cell development. To test the role of FAM72A in CSR in primary B cells, we isolated splenic B cells from *Fam72a*^−/−^ and *Fam72*^+/+^ mice and performed an *ex vivo* CSR assay. We found that switching to all Ig isotypes was defective in *Fam72a*^−/−^ compared to *Fam72*^+/+^ mice (**Fig. 2a**). *Fam72*^+/−^ mice and *Fam72*^+/−^ CH12 cells had a partial defect in CSR (**Extended data Fig. 6a,b**) suggesting that FAM72A is haploinsufficient. We immunized mice with NP-CGG and found that the generation of IgG^+^ NP-specific antibody-secreting cells was reduced in *Fam72a*^−/−^ compared to *Fam72*^+/+^ mice (**Fig. 2b**). FAM72A-deficiency did not impact the expression of *Aicda* and *Ig* germ line transcripts, cell proliferation, or cell cycle progression in B cells (**Extended data Fig. 6c-f**). These data indicate that FAM72A is critical for CSR in primary mouse B cells.

**Figure 2.**
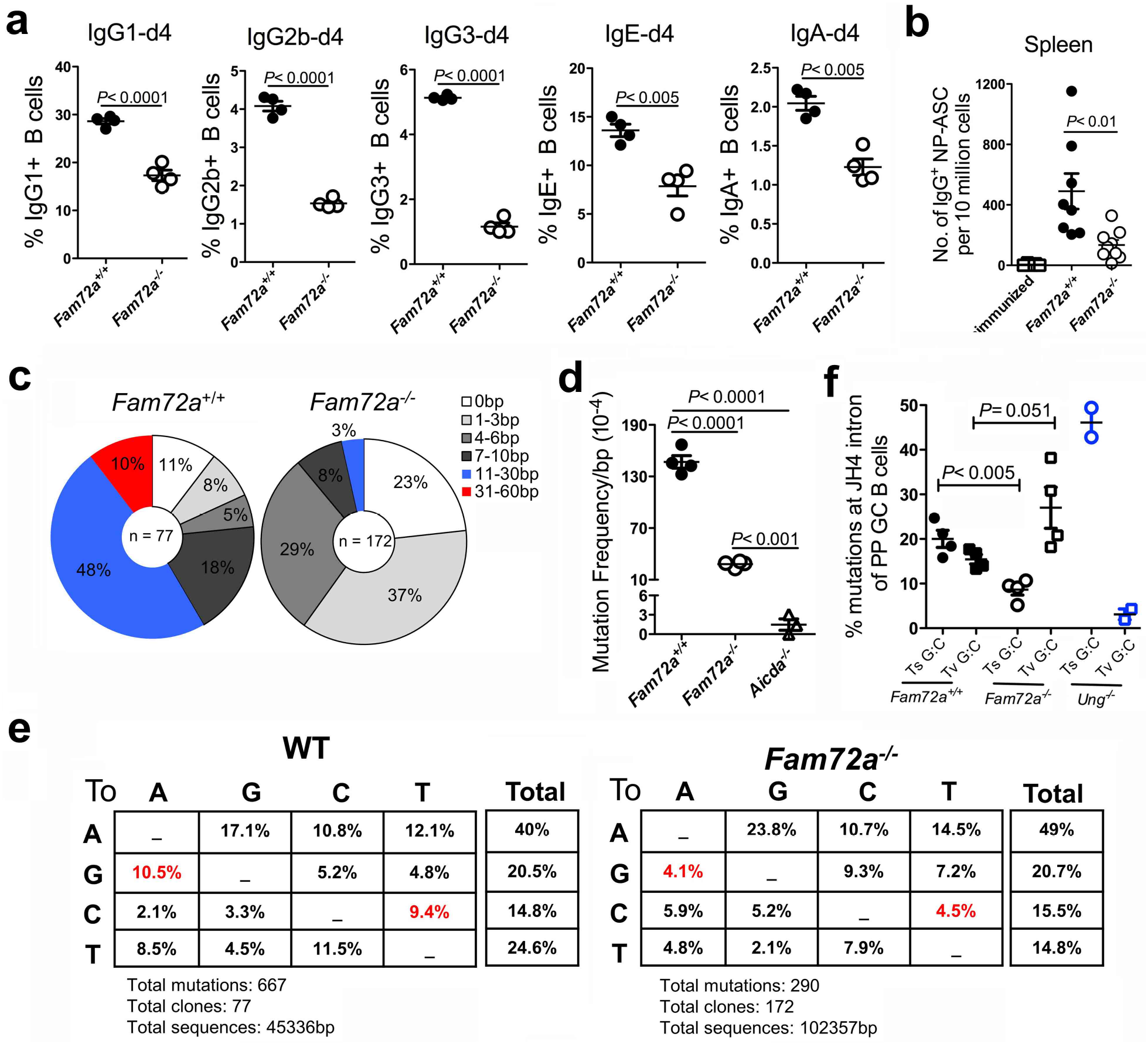
*Fam72a*^−/−^ mice exhibited defects in CSR and somatic hypermutation. (a)Analysis of *ex vivo* CSR to indicated isotypes using splenic B cells from *Fam72a*^−/−^ or *Fam72a*^+/+^ littermate mice (n= 4 mice per group). Data are representative of 2 independent experiments. **(b)** Enumeration of IgG^+^ NP-specific antibody secreting cells (ASC) from NP29-CGG immunized mice or unimmunized control (n= 5-9 mice per group; data were pooled from 2 independent experiments). **(c-e)** Analysis of somatic hypermutation profiles in the JH4 region of germinal center B cells that were sorted from Peyer’s patches of naïve 5-month old *Fam72a*^−/−^ or *Fam72a*^+/+^ littermate mice (n= 4 mice per group). The total amount of sequences analyzed (center of pie charts) and proportions of sequences with indicated numbers of mutations are shown in (**c**). The frequency of unique mutations in the JH4 region of germinal center B cells per mouse was shown in (**d**) (n=3 mice for *Aicda*^−/−^ group). (**e**) The compiled spectrum of unique top-strand (coding-strand) mutations. These two panels represented the pooled data from 4 *Fam72a*^+/+^ and 4 *Fam72a*^−/−^ mice, respectively. The mutation spectrum for each individual mouse is shown in **Extended data Fig 8**. (**f**) The frequencies of transitional (Ts) and transversional (Tv) mutations at G:C pairs in the JH4 region of germinal center B cells from Peyer’s patches of naïve *Fam72a*^−/−^ or *Fam72a*^+/+^ littermate mice (n= 4 mice per group). Transitional G:C mutations include G->A and C->T mutations, while transversional G:C mutations include G->C or T, and C->A or G mutations. The data using the *Ung*^−/−^ mice (n=2 mice) was obtained from ^9^. Data in (**a**, **b**, and **d**) were presented as mean ± SEM and were analyzed using two-tailed unpaired Student t test.

To explore a role of FAM72A in SHM, we isolated germinal center B cells from Peyer’s patches of *Fam72a*^−/−^ and *Fam72*^+/+^ mice (*Aicda*^−/−^ mice were used as background mutation control), and sequenced the JH4 region (**Extended data Fig. 7a**). The proportion of GC B cells in Peyer’s patches was comparable between *Fam72a*^−/−^ and *Fam72*^+/+^ mice (**Extended data Fig. 7b**). However, the loss of *Fam72a* caused a ~4 fold reduction in mutation frequency in the JH4 region in GC B cells (**Fig. 2c,d**, and **Extended data Fig. 8**). While all C:G and A:T mutations in the JH4 region were reduced in the *Fam72a*^−/−^ mice, the characteristics of the remaining mutations were altered in the *Fam72a*^−/−^ mice (**Fig. 2e**): B cells from *Fam72a*^−/−^ mice had a reduced frequency of transition mutations at G:C basepairs and an increase in transversion mutations at G:C basepairs (**Fig. 2e**). Strikingly, the transition versus transversion mutation preferences at G:C basepairs in *Fam72a*^−/−^ mice was opposite to that found in *Ung*^−/−^ mice (**Fig. 2f**) ^9^ and more in line with overexpression of UNG2 ^17^. Overexpression of another uracil DNA glycosylase, SMUG1, has also been shown to reduce SHM in mice ^18^. These data suggest that UNG2 is hyperactive in the absence of FAM72A.

CSR completion depends on two double-stranded DNA break (DSB) repair pathways that perform the end joining of the DSBs ^2,19–22^. To explore the underlying mechanism behind the effect of FAM72A on CSR, we asked whether FAM72A functions in non-homologous end joining (NHEJ) and alternative end joining (A-EJ). Using DNA-repair specific substrates with GFP expression as a read out, we found that NHEJ and A-EJ were unaffected in FAM72A-deficient CH12 clones (**Extended data Fig. 9**). We then examined in *Fam72a*^−/−^ CH12 clones the mutations 5’ of the μ switch region, which reflect a balance between the mutagenic activity of AID during transcription, as well as faithful repair by BER and MMR ^8^. We found that the mutation frequency was reduced in *Fam72*^−/−^ CH12 clones compared to controls (**Fig. 3a, Extended data Fig. 10**). The reduced mutation frequency at the μ switch region was not due to reduced AID expression or altered μ germ line transcripts in *Fam72*^−/−^ CH12 clones (**Fig. 3b,c**). As the reduced mutation frequency could be due to altered dU processing in FAM72A-deficient cells, we tested whether FAM72A impacted AID-induced mutations in the *Ung*^−/−^ background. We generated *Ung*^−/−^ and *Ung*^−/−^ *Fam2a*^−/−^ CH12 clones (**Extended data Fig. 2b**). As would be expected, the mutation frequency 5’ of the μ switch region was elevated ~3-fold in the *Ung*^−/−^ CH12 clone compared to WT controls owing to the lack of repair of AID-induced dU (**Fig. 3a**). Importantly, the loss of *Fam72a* in the *Ung*^−/−^ background did not reduce the mutation frequency (**Fig. 3a**). These data suggest that AID activity was unaffected by FAM72A-deficiency, and that FAM72A affects SHM through UNG2.

**Figure 3.**
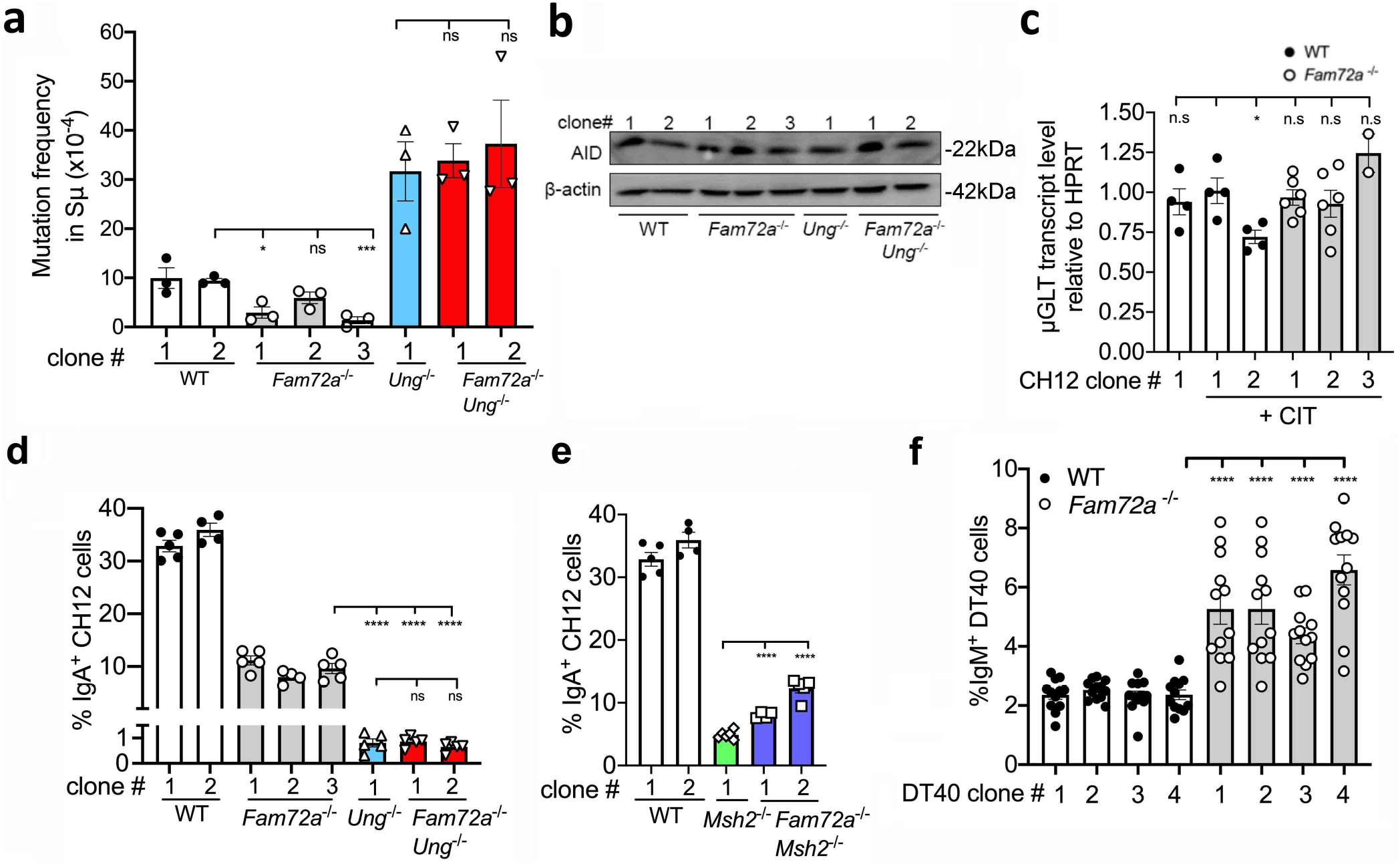
FAM72A is epistatic with UNG during CSR. (**a**) Sμ mutation frequency in WT, *Fam72a*^−/−^, *Ung*^−/−^, and *Fam72a*^−/−^*Ung*^−/−^ CH12 cells. CH12 cells were treated with CIT for 5 days, followed by DNA extraction and PCR amplification of the region 5’ of Sμ. The mutation frequency was calculated by counting the number of unique mutations divided by the total number of nucleotides sequenced. Data are representative from 3 independent experiments. (**b**) AID expression levels of WT, *Fam72a*^−/−^, *Ung*^−/−^, and *Fam72a*^−/−^*Ung*^−/−^ CH12 cells as assessed by Western blot analysis. β-actin was used as loading control. (**c**) Sμ germ line transcripts (μGLT) of the indicated CH12 clones before and after stimulation with CIT for 2 days. (**d**) CSR was assessed in WT, *Fam72a*^−/−^, *Ung*^−/−^, and *Fam72a*^−/−^*Ung*^−/−^ clones. Cells were treated with CIT for 2 days, followed by flow cytometry analysis for surface IgA expression. *, p<0.05; **, p<0.01; ***, p<0.001; ****, p<0.0001. (**e**) Same as d, except that CSR was analyzed in WT, *Msh2^−/−^*, and *Fam72a*^−/−^*Msh2*^−/−^ CH12 clones. Data shown in panels (d) and (e) were performed at the same time. (**f**) Fluctuation analysis of gene conversion in WT and *Fam72a*^−/−^ DT40 clones at two weeks of expansion.

To further test the notion that FAM72A functions with UNG2, we carried out CSR assays in cells doubly deficient in FAM72A and UNG, as well as those doubly deficient in FAM72A and MSH2 (**Extended data Fig. 2b, c**). *Ung*^−/−^ CH12 cells switched at ~3% of WT CH12 cells (**Fig. 3d**). However, knocking out *Fam72a* in UNG-deficient CH12 cells did not further reduce CSR (**Fig. 3d**), suggesting that FAM72A is epistatic with UNG2 during CSR. On the other hand, *Msh2*^−/−^ CH12 cells switched at 15% of WT CH12 cells (**Fig. 3e**). Notably, the effects on CSR of knocking out either *Ung* or *Msh2* were greater than 85% and therefore not additive, suggesting that like for SHM ^11^, the concerted action of UNG and MMR pathways promotes CSR. Strikingly, knocking out *Fam72a* in MSH2-deficient CH12 cells led to a modest, but significant increase in CSR (**Fig. 3e**). This result is consistent with FAM72A inhibiting UNG2 since in the MSH2-deficient background, the only sources of DNA breaks would derive from UNG2, and increased UNG2 activity would lead to increased breaks necessary for CSR.

To further examine the relationship between FAM72A, UNG2, and mismatch repair, we tested whether FAM72A modulates *Ig* gene conversion, another antibody diversification process initiated by AID ^23^. In this case, gene conversion is catalyzed by UNG2 ^24^ but not by mismatch repair ^25^. We generated *Fam72a*^−/−^ DT40 clones (**Extended data Fig. 11a**) and found that gene conversion was increased ~2-fold relative to controls (**Fig. 3f**). *Fam72a*^−/−^ DT40 clones had a reduced proliferation rate than controls (**Extended data Fig. 11b**) and Ig gene conversion increases linearly with cell generations ^25^. Nonetheless, analysis of the data using the same number of cellular divisions showed the same effect of increased gene conversion in *Fam72a*^−/−^ DT40 cells (**Extended data Fig. 11c**). The result is conceptually similar to the effect on CSR after knocking out *Fam72a* in *Msh2*^−/−^ CH12 cells. Notably, chicken MSH2/6 fails to recognize U:G mismatches in DT40 cells ^25^, so UNG would have no competition from faithful MMR in Fam72a-deficient DT40 cells, explaining the increase in Ig gene conversion. Collectively, these data support the notion that FAM72A modulates 3 antibody diversification processes by limiting UNG2.

A previous report suggested that FAM72A interacts with UNG2 ^16^. To confirm this interaction, we performed a BioID proximity-labeling assay and found that human FAM72A is in close proximity to UNG in HEK-293 cells (**Fig. 4a**). To test a direct interaction, purified murine versions of these proteins (6xHis-mFAM72A and mUNG2-FLAG, **Extended data Fig. 12a**) were mixed at 1:1 ratios, and the pull-down of the proteins using either nickel-containing beads or anti-FLAG antibodies resulted in co-precipitation of the other protein (**Fig. 4b**). This interaction was conserved across mammalian species as pull-down of mFAM72A resulted in co-precipitation of hUNG2 (**Fig. 4b**).

**Figure 4.**
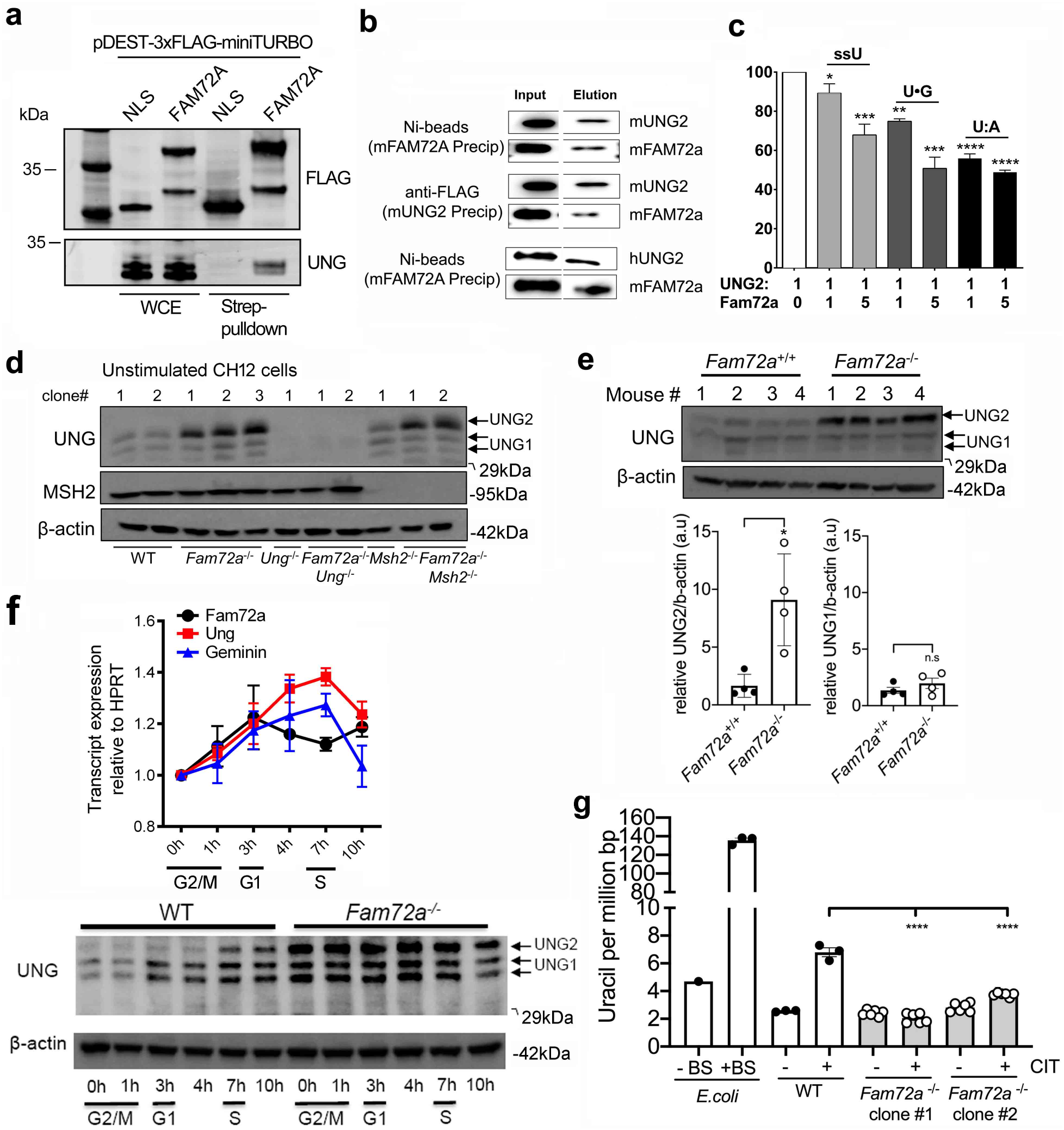
FAM72A binds to and inhibits UNG2 and leads to reduced UNG2 protein levels. (**a**) Streptavidin affinity purification from 293T cell lysates expressing N-terminal 3xFLAG-miniTURBO tagged FAM72A upon proximity biotinylation. FLAG-miniTURBO-SV40-NLS was used as control. Bound proteins and whole-cell extracts (WCEs) were immunoblotted with the indicated antibodies. (**b**) Murine UNG2 and FAM72A proteins were mixed in equal molar amounts and pulled-down using polyHis-tag or FLAG-tag on mFAM72A and mUNG2 proteins, respectively. The input and elutions were analyzed by Western blot using anti-His tag or anti-FLAG tag antibodies. Cross-species pull-down of mFAM72A with hUNG2 is also shown: the hUNG2 protein was detected using anti-UDG antibody. (**c**) Purified murine UNG2 and Fam72A proteins were pre-incubated then reacted with either a single-stranded uracil substrate (ssU) or double-stranded uracil substrates with either U:G or U:A pairs. Quantification of three independent experiments of the type shown in **Extended data Fig. 12** busing 0:1, 1:1 or 5:1 molar ratios of mFAM72A to mUNG2. (**d**) Western blots for UNG, MSH2, and β-actin (as control) in unstimulated CH12 clones of the indicated genotype. UNG1 and UNG2 are indicated on the gel. (**e**) Western blots for UNG and β-ACTIN in primary B cells isolated from WT and *Fam72a*^−/−^ mice. Quantitation of multiple blots shown below shows increased UNG2 but not UNG1 in *Fam72a*^−/−^ B cells. (**f**) *Fam72a*^+/+^ CH12 cells were synchronized at G2/M phase for 20h with RO-3306 treatment, and then cells were harvested at 0h (G2/M), 1h (G2/M), 3h (G1), 4h, 7h (S), and 10h post RO-3306 release, followed by qPCR analysis of *Fam72a*, *Ung* or *Geminin* (accumulating at S phase) mRNA (top panel). In bottom panel, cells were harvested, and lysates were examined by western blot for UNG and β-ACTIN protein by western blot. **(g)** Quantification of genomic uracils in DNA isolated from WT and *Fam72a*^−/−^ CH12 clones 1 and 2 was performed using the method described in ^36^. The quantification was done in technical triplicates.

To test whether FAM72A modulates UNG2 activity, we carried out *in vitro* uracil excision assays. The addition of mFAM72A to the mUNG2 reaction resulted in a moderate reduction in the ability of UNG2 to excise uracils from three different types of substrates: single-stranded DNA with a uracil, double-stranded DNA with a U•G mispair or a U•A pair (**Fig. 4c, Extended data Fig. 12b**). mFAM72A also moderately inhibited human UNG2, but had no effect on the activity on *E. coli* UNG (**Extended data Fig. 12c**). However, FAM72A had a striking effect on UNG2 protein levels, as *Fam72a*^−/−^ CH12 cells had significantly increased UNG2 but not UNG1 protein levels (**Fig. 4d, Extended data Fig. 12e**). This result was confirmed in primary B cells from *Fam72a*^−/−^ mice (**Fig 4e**). These data suggest that FAM72A binds to and antagonizes UNG2 in two different manners.

To test whether the expression of *Fam72a* is cell cycle-dependent, we synchronized CH12 cells by arresting them at G2-M using the CDK1/cyclin B1 inhibitor RO-3306, and then released the blockade. *Ung* and *Germinin* levels peaked at 7 hours after the inhibitor was removed (**Fig. 4f**), suggesting entry into S-phase. However, *Fam72a* transcripts peaked at 3 hours after the inhibitor was removed (**Fig. 4f**) suggesting that *Fam72a* is elevated in G1, and might impact UNG2 levels in G1. Indeed, probing UNG protein by western blots in these synchronized cells shows that UNG2 protein is markedly increased in the G1 phase of the cell cycle in *Fam72a*^−/−^ CH12 cells compared to controls (**Fig. 4f**, **Extended data Fig. 12f**), a stage of the cell cycle that coincides with AID-induced deaminations ^26^. The increased UNG2 levels in *Fam72a*^−/−^ CH12 affects the genome-wide uracil content following CIT stimulation, as we observed that the amount of genomic uracils more than doubled following CIT stimulation and this increase was completely eliminated in *Fam72a*^−/−^ CH12 clones (**Fig. 4g**). These data together show that FAM72A limits UNG2 in cells allowing the accumulation of dUs in U:A or U•G pairs following AID-mediated deamination of deoxycytidines in DNA. While these excessive dUs have long been proposed to cause SHM through replication and error-prone base excision repair (BER), the U•G mispairs are also the trigger for aberrant MMR which contributes to CSR (**Extended data Fig. 13**). Thus, FAM72A plays a key role in creating the raw material for both SHM and CSR and connects the BER and MMR processes.

The key limitation of using AID-induced dUs for SHM and CSR, is that mammalian UNG2 is abundant in the nucleus ^27^ and has a high turnover rate ^28^. Such a high uracil excision activity is required because dUTP is present in the nuclei at appreciable concentrations ^29^ and replicative DNA polymerases readily misincorporate this non-canonical base into DNA. UNG2 efficiently replaces dU with dT through BER keeping the level of dU in DNA at ~1 in 10^6^ bp ^30^ or below ^31^. Consequently, any dUs created by AID are likely to be quickly excised by UNG2 eliminating U•G mispairs that are required for the involvement of MMR in SHM and CSR.

FAM72A alleviates this problem largely by reducing UNG2 protein levels and its activity in B cells. FAM72A therefore affects SHM in two ways. First, by decreasing uracil excision, it increases the proportion of U•G mispairs that can persist until replication thereby creating C:G to T:A mutations. Second, it engages MMR by increasing U•G mispairs, leading to the recruitment of POLη and the introduction of mutations at A:T base pairs^32^ (**Extended data Fig. 13**). Thus, the presence of FAM72A increases SHM frequency both at C:G and A:T pairs, as observed in our study. The increased U•G mispairs caused by FAM72A at switch regions would also lead to increased engagement of MMR to promote CSR. UNG2 protein levels are regulated by cell cycle dependent phosphorylation ^33^, and potentially provides a mechanism for the regulation of UNG2 protein levels by FAM72A.

FAM72A is found predominantly in jawed vertebrates, but is also found in other eukaryotes, including some fungal species. This suggests that, albeit playing a principal role in modulating the antibody immune response, FAM72A might have other ancestral functions, such as in DNA repair. Of particular interest is the presence of 4 almost identical paralogues of FAM72 in humans, termed FAM72A-D ^34,35^, that are not present in other vertebrates, including non-human primates. Whether these human paralogues play a role similar to mFAM72A in modulating antibody maturation warrants further studies.

## Methods

### Mice

*Fam72a* mutant sperm: C57BL/6N-Fam72a^tm1.1(KOMP)Vlcg^/MbpMmucd, in which a 10,298bp region between 131, 528,704 to 131, 539, 001 positions of mouse chromosome 1 was deleted by the insertion of ZEN-Ub1 cassette, was purchased from Mutant Mouse Resource and Research Centers (UC Davis, CA). The *in vitro* fertilization of *Fam72a* mutant sperm with C57BL/6N female mice (Charles River) was performed at The Centre for Phenogenomics (Toronto, Canada). *Fam72a* heterozygous mice were bred in our animal facility and maintained under pathogen-free conditions. The experimental procedures were approved by the Animal Care Committee of University of Toronto.

### Plasmids

For Cas9-mediated gene editing, gRNAs were cloned into px330 (Addgene, plasmid #42230), lentiGuide-Puro (Addgene, plasmid #52963) or lentiCRISPR v2 (Addgene, plasmid #52961). Murine *Fam72a* was PCR amplified from CH12 cDNA and cloned into a pRSET-A vector (Thermo Fisher). The murine UNG2 (mUNG2) was amplified from mUNG2 cDNA (OriGene), and cloned into FLAG-HA-pcDNA3.1 vector (Addgene, plasmid #52535) using Gibson assembly kit (New England Biolabs) to create pcDNA3.1-mUNG2-polyGly-FLAG (mUNG2-FLAG) expression construct. The mUNG2-FLAG gene was amplified and cloned into a pET28a^+^ expression vector. The coding sequence for human FAM72A (hFAM72A) was ordered from IDT with gateway compatible flanking sequences and cloned into pDEST-NLS-3xFLAG-miniTURBO (gift from Dr. Anne-Claude Gingras laboratory, LTRI, Toronto) via gateway cloning. The HR substrate (DR.GFP) is a gift from Dr. Maria Jason, NHEJ substrate (EJ5-GFP) is a gift from Dr. Takashi Kohno, and A-EJ substrate (EJ2-GFP) is a gift from Dr. Jeremy Stark. All the plasmid constructs were verified by Sanger sequencing. Primers were obtained from Invitrogen and are listed in Extended data Table 1.

### Lentiviral transduction

To generate lentiviruses, HEK293T cells were seeded in DMEM with 10% FBS and Penicillin-Streptomycin, and 18 h later, were transfected with lentiviral packing plasmids pMD2.G (Addgene, plasmid #12259) and psPAX (Addgene, plasmid #12260) along with lentiviral plasmids containing gRNAs of interest in the presence of PEI. The lentivirus-containing supernatants were harvested at 48h to 96h post-transfection and concentrated by ultracentrifugation. The lentiviral transduction of CH12 cells were performed via centrifugation at 800xg for 1h at room temperature in the presence of 8μg/ml polybrene.

### CRISPR-Cas9 screen in CH12F3-2 cell line

CH12 cells were cultured as we previously described ^1^. To generate Cas9-expressing CH12 cells, CH12 cells were transduced with LentiCas9-Blast (Addgene, plasmid #52962) and selected in the presence of 10 μg/ml Blastcidin for 9 days. Single clones that stably expressed high levels of Cas9 were collected to generate the pooled Cas9-expressing CH12 cells. The timeline that allowed maximal Cas9 editing efficiency in CH12 cells was determined by monitoring CSR defects caused by gRNAs that target AID. Seventy-one million IgA^−^ CH12 cells stably expressing Cas9 were transduced with the lentiviral-based mTKO library ^2^ consists of 95,528 gRNA sequencs (pLCKO-mTKO) at a MOI of 0.6, followed by puromycin selection 2 days. The pLCKO-mTKO library is available at Addgene (https://www.addgene.org/pooled-library/moffat-mouse-knockout-mtko/). CH12 cells were subsequently stimulated with recombinant TGFβ (R&D systems), recombinant IL-4 (R&D systems), and anti-CD40 (ThermoFisher) for 3 days, as we previously described (PMID: 27160905). The cells were stained for surface expression of IgA using PE-conjugated anti-mouse IgA antibody (SouthernBiotech) and sorted by a FACS Aria IIIu and influx sorter (BD Biosciences). The CRISPR sequencing library was prepared as described before ^3^. Genomic DNA from T0 and T9 (IgA+, IgA- and unsorted) samples was extracted using the Wizard Genomic DNA Purification kit (Promega) according to manufacturer’s instructions. To generate sequencing libraries, gRNA barcode sequences were amplified as described previously ^2^. The resulting libraries were sequenced on an Illumina HiSeq2500. Raw and normalized read counts will be deposited in NIH BioProjects upon publication.

### CRISPR/Cas9-mediated gene editing in CH12 cells and DT40 cells

To knock out gene of interest in CH12 cells, the px330 vector containing gRNA against *Fam72a, Ung* or *Msh2* was electroporated into CH12 cells. At 72h post-electroporation, CH12 cells were subcloned by limiting dilution. Three *Fam72a*^−/−^ clones were generated (12A, 2B and 6H) and for the purpose of simplicity, were renamed as clone #1, 2 and 3, respectively, while two *Fam72a*^+/+^ clones (20C and 23C, renamed as clone #1 and 2) were used as control. Individual knockout clones were sequenced and validated using qPCR and/or Western blot. DT40 CL18 *Fam72a*^−/−^ cells were obtained by transfecting DT40 CL18 with Nucleofector™ Kit V four constructs in PX458 (Addgene, plasmid #48138) targeting *Fam72a* (1.5μg each) and two pBluescript KS(+) constructs containing an antibiotic cassette resistance each (either for Blasticidin or Hygromycin) flanked by *Fam72a*-homology arms (2μg each). At 48h post-electroporation, Blasticidin (25μg/mL) and Hygromycin (2mg/ml) were added and cells were plated in 96-well plates (6000 cells in 20μL/well). After 7–14 days, cells from wells containing a single colony were expanded. Genomic DNA was isolated and knock out clones were identified by PCR and verified by RNA extraction and RT-PCR (Extended data Fig 11a). All gRNA sequences are listed in Extended data Table 1.

### Measuring expression of *Fam72a* and *Ung* mRNA at different stages of the cell cycle

CH12 cells were synchronized at G2/M phase, as previously described ^4^. In brief, *Fam72a*^+/+^ CH12 cells (20 × 10^4^ cells/mL) were cultured 20 hours with 10uM of CDK1/cyclin B1 inhibitor RO-3306 (Sigma), followed by exchange into inhibitor-free culture media. Cells were harvested at 0h (G2/M), 1h (G2/M), 3h (G1), 4h, 7h (S), and 10h post-RO-3306 release, followed by RNA preparation and qPCR analysis of *Fam72a* and *Ung* mRNA. The corresponding cell cycle of CH12 cells at these collection timepoints were previously confirmed by ^4^ in CH12 cells using DNA staining with propidium iodide (detected by flow cytometry), which was validated by qPCR analysis of *Geminin* mRNA, which accumulates in S phase.

### Monitoring growth and Ig Gene Conversion in DT40 cells

To determine DT40 cells doubling time, cultures of 5×10^4^ cells/mL per condition were set and counted in duplicate by hemocytometer every 6 hours for 4 days. Data was analyzed with GraphPad Prism and doubling time was determined by nonlinear regression of data to Exponential (Malthusian) growth equation (Extended data Fig 11b). Immunoglobulin Gene Conversion was monitored by the conversion of IgM- to IgM+ cells by flow cytometry. Fluctuation analysis was performed as described ^5^ with modifications. Briefly, 4 single cell clones of DT40 WT or Fam72a^−/−^ were stained with anti-IgM-PE (1:40, clone M-1, SouthernBiotech) and 12 populations of 5×10^4^ IgM^−^ cells were sorted for each clone. The proportion of IgM-gain was measured by flow cytometry using the same staining after 2 weeks of expansion (same time of expansion) and 5 days later for the Fam72a^−/−^ cells (to achieve same number of divisions than the wt at two weeks). Data was analyzed with FlowJo™ and GraphPad Prism.

### Human FAM72A proximity-dependent BioID

HEK293T were seeded in 10cm dishes, and 24 h later cells were transfected by standard PEI method with pDEST-NLS-3xFLAG-miniTURBO (1.5 μg) or pDEST-NLS-3×FLAG-miniTURBO-FAM72A (3 μg). At 48 h post-transfection, 5 μgr/mL doxycycline was added and 24 h later, cells were exposed to Biotin (50 μM) for 1 h. Cells were washed with PBS, scraped and pelleted. Cell pellet was lysed in 0.5 mL of lysis buffer (50 mM Tris-HCl pH 8.0, 100 mM NaCl, 2 mM EDTA, 0.5%, NP-40 and 10 mM NaF, 10 mM MgCl2) and 250 U of Benzonase per sample on ice for 15 min. SDS was added to a final concentration of 0.4 % and incubated for 5 min at 4° C. Lysates were centrifuged at 15,000xg for 5 min and supernatants were incubated with 50 μL of Streptavidin Sepharose High Performance beads (GE Healthcare) for 1 h rotating. Beads were washed 5 times with lysis buffer and bound proteins were eluted by boiling 5 min at 96° C in SDS-PAGE sample buffer and analyzed by immunoblotting.

### Primary B cell analysis

B cell development in bone marrow and spleen of sex-matched *Fam72a*^−/−^ and *Fam72a*^+/+^ littermate (6-8 weeks old) was evaluated, as we previously described ^6^. To induce *ex vivo* CSR to different Ig isotypes, splenic B cells were purified with a mouse B cell isolation kit (StemCell Technologies), and then stimulated with lipopolysaccharide (LPS; Sigma) in combination with various cytokines, as we previously described ^6^. Cells were collected at day 4 post-stimulation and were stained with antibodies against mouse IgG1 (PE, BD Pharmigen), IgG2b (PE, SouthernBiotech), IgG3 (FITC, BD Pharmigen), IgE (FITC, BD Pharmigen), or IgA (PE) to assess CSR. For the cell cycle analysis, splenic B cells were stimulated with LPS for 2.5 days, before the Click-iT Plus EdU Alexa Fluor 647 kit (ThermoFisher Scientific) was utilized, as we previously described ^6^. The flow cytometric data were analyzed with a FlowJo X10 software.

### Analysis of Sμ and JH4 region mutations

For Sμ region sequencing in CH12 cells, wild-type, *Fam72a*^−/−^, *Ung*^−/−^, and *Fam72a*^−/−^*Ung*^−/−^ CH12 cells that have been cultured with CIT for 5 days. Corresponding genomic sequence from resting wild-type CH12 cells were used as the reference sequence. For JH4 region mutation analysis, Peyer’s patches were harvested from approximately 5-month old *Fam72a*^−/−^, *Fam72a*^+/+^ littermate, and *Aicda*^−/−^ mice, respectively. Single cell suspensions of Peyer’s patches were prepared via grinding, and then filtered through a 70-μm cell strainer. As we previously described ^7^, cells were stained with antibodies against mouse B220 (PE, clone: RA3-6B2, eBiosciences), Fas (biotin, clone: Jo2, BD Pharmigen) and GL-7 (eF660, clone: GL-7, eBiosciences) in the presence of Fc block (2.4G2), followed by streptavidin-APC-eFluor 780 staining, to identify germinal centre B cells. The cells were then stained with 7-AAD (live/dead marker) prior to flow cytometric sorting on an FACS Aria II sorter (BD Biosciences). The sorted germinal centre B cells (7-AAD^−^/B220^+^/GL-7^+^/Fas^+^) were digested with Proteinase K (Invitrogen) at 55°C overnight, followed by phenol-chloroform genomic DNA purification. The JH4 intron region was PCR amplified using a high-fidelity Platinum SuperFi II DNA polymerase (Invitrogen) with primers listed in Extended data Table 1. Gel-extracted JH4 PCR products were cloned and sequenced at cloned into Zero Blunt vectors (Invitrogen) or Clone JET (Invitrogen) and then transformed into competent DH5α bacteria. The plasmids yielded from individual bacterial clones were sequenced at The Centre for Applied Genomics (Toronto, ON). Sequences were analyzed with DNA-STAR software and were aligned with the corresponding genomic sequences as described ^8^.

### Quantitative real-time PCR

RNA extraction was performed using TRIzol (Invitrogen), followed by DNase I treatment (Invitrogen), and reverse transcription with Maxima First strand cDNA synthesis Kit for real-time PCR (Thermo Fisher) to prepare cDNA. The cDNA samples were subjected to quantitative PCR (qPCR) reactions using PowerUp™ SYBR Green Master Mix (Applied Biosystems) according to the manufacturers protocol. Primer sequences are listed in Extended data Table 1.

### Purification of murine UNG2 and FAM72A proteins

The plasmid pRSET_6XHis-mFAM72A was transformed into BL21(DE3) and mFAM72A expression was induced using 0.1 mM IPTG and the cells were grown at 18°C for additional 18 hours. The cells were harvested by centrifugation and resuspended in 30 mL of lysis buffer (20 mM Tris-Cl pH 8.0, 50 mM NaCl, 10 mM imidazole) supplemented with cOmplete™ protease inhibitor cocktail. Cells were lysed using a French Press, the lysate clarified using centrifugation. The supernatant was mixed with the Ni-NTA agarose beads and rotated at 4°C for 1 hour. The beads were successively washed with 20 mL of Buffer A (20 mM Tris-Cl pH 8, 50 mM imidazole) containing 50, 250, or 500 mM NaCl. The mFam72A protein was eluted from the beads using 500 μL aliquots of elution buffer (20 mM Tris-Cl pH 8.0, 50 mM NaCl, 250 mM imidazole). The elution fractions found to have the protein when analyzed on an SDS-PAGE gel were combined and dialyzed using Buffer B (20 mM Tris-Cl pH 8.0, 50 mM NaCl, 1 mM DTT, 1 mM EDTA, 10% glycerol). The protein was concentrated to 1 mg/mL using 10kDa MWCO centrifugal filter and stored at −80°C.

The BL21(DE3) cells containing plasmid pET28a-mUNG2-6XGly-FLAG were grown, the protein expression was induced using IPTG (0.5 mM) and the cells were broken in the same way as cells expressing mFam72A (see above). Following cell lysis, the cleared lysate was mixed with the anti-FLAG M2 magnetic beads (SIGMA-ALDRICH) and rotated at 4 °C for 18 hours. The beads were harvested using a DynaMag-2 magnet (Thermofisher) and successively washed with 1XTBS-T containing 50, 250, or 500 mM NaCl. The mUNG2 protein was eluted with 500 μL of FLAG Elution Buffer (3XFLAG peptide at 150 ng/μL in 1XTBS). The elution fractions found to have the protein when analyzed on an SDS-PAGE gel were combined, the buffer was exchanged 3 times with UNG2 storage buffer (25 mM HEPES–NaOH (pH 7.4), 200 mM NaCl, 0.01% Triton X-100 and 1 mM TCEP, 20% glycerol) and the protein was concentrated to 5 mg/mL using 10kDa MWCO centrifugal filter and stored at −80°C. The human UNG2 protein was a generous gift from Dr. Brian Weiser (Rowan University School of Osteopathic Medicine).

### Co-precipitation of mFAM72A and mUNG2 proteins

The UNG2 proteins were mixed with mFAM72A at a molar ratio of 1:1 in the UNG2 buffer and incubated at 25°C for 20 minutes. The proteins were co-precipitated by respectively pulling down mUNG2-FLAG or 6XHis-mFam72A using anti-FLAG magnetic beads or Ni-NTA agarose beads, respectively. After an overnight incubation at 4°C while rotating, the beads were washed 3 times with 1XTBS-T and eluted by the addition of Laemmli buffer and boiling the mixture for 10 minutes at 95°C. The input and elution samples were analyzed by Western blot by probing with mouse anti-FLAG M2 antibody (Sigma Aldrich; 1:2000), rabbit anti-His tag antibody (Cell signaling; 1:2000), and anti-UDG antibody (Santa Cruz; 1:500) followed by the addition of goat anti-rabbit IgG and goat anti-mouse IgG HRP-conjugated antibodies (Cell signaling; 1:1000). The protein bands were visualized by the addition of Super signal West Pico Plus chemiluminescence substrate (ThermoFisher) and detected using a FluorChemQ scanner (Cell Biosciences Inc.).

### In vitro uracil excision assay

DNA oligomers containing uracil used in these assays are listed in the Extended data Table 1. The mUNG2 protein alone or mixed with mFAM72A was preincubated at 25°C for 20 minutes in UNG2 buffer (10 mM Tris–HCl (pH 8.0), 100 mM NaCl, 1 mM DTT and 5 mM EDTA). This was followed by incubation at 25°C for 5 minutes and the reactions were stopped by the addition of NaOH to 0.1 M and boiling at 95°C for 10 minutes. Formamide dye was added to 50% v/v and the reaction products were separated on a 15% denaturing polyacrylamide gel. The DNA was visualized by scanning for Cy2 fluorescence using a Typhoon FLA 9500 phosphor imager and band intensities were quantified using ImageJ software. The percent product was normalized to the negative control sample (mUNG2 without mFAM72A) and bar graphs were generated using GraphPad Prism 8 and the statistical significance determined using Mann-Whitney U test at a 95% confidence interval.

### Quantification of genomic uracils

The genomic uracils were analyzed in CH12 cells as described previously ^9^. Briefly, about 5 μg of the genomic DNA was digested with HaeIII (New England Biolabs) and purified using phenol:chloroform extraction followed by ethanol precipitation. The digested DNA was incubated with AA7 (Sigma Aldrich, 10 mM final concentration) at 37 °C for 1 hour to block pre-existing abasic sites. DNA was then treated with

*E. coli* uracil DNA-glycosylase at 37 °C for 30 minutes followed by 1-hour incubation with 2 mM AA6. AA6 tagged DNA was labeled with DBCO-Cy5 (1.7 μM) by shaking the reaction mixture for 2 hours at 37 °C in dark. The tagged DNA was purified using DNA clean and concentrator kit (Zymo research). Fluorescently labeled DNA was transferred on to Zeta probe membrane (Bio Rad) using a dot blot apparatus (Bio-Rad) and the membrane was scanned using a Typhoon 9210 phosphor imager (GE Healthcare). The images were analyzed using the ImageJ software. Fluorescence intensity was converted to number of uracils using the CJ236 standard curve as described previously^9^.

### Western blotting

Western blotting was performed as we previously described ^10^. The following antibodies were used: anti-UNG (abcam; cat# 245630), anti-AID (Invitrogen; cat# 39-2500), anti-MSH2 (Invitrogen; cat# 33-7900), or β-ACTIN (Sigma; cat# A2066). Densitometry analysis was performed using Image J (NIH).

## Statistical Analysis

All analyses were performed on GraphPad Prism software. Otherwise noted, two-tailed unpaired student t test was used.

## Data availability

All relevant data are available from the authors

## Acknowledgements

We thank Dr. Philippe Poussier for critical review of the manuscript. YF, CL, and AAQ are recipients of the Canadian Institutes of Health Research Postdoctoral Fellowship. NSD is the recipient of a doctoral award from the FRQ-S (Fonds de recherche Santé Québec). JAS was supported by a competitive graduate research assistantship from the Wayne State University and RM-R was supported by a Thomas C. Rumble fellowship. JMDN is a Merit scholar from the Fonds de recherche de Quebec – Santé, and is supported by CIHR (PJT-155944). JM is supported from a CIHR project grant (CBT-438323) and holds a Canada Research Chair in Functional Genomics. DD is a Canada Research Chair (Tier I) and work in the DD lab was supported by a grant from the CIHR (FDN143343). ASB was supported by a National Institutes of Health grant (1R21AI144708) and Bridge Funding grant from Wayne State University. AM is supported by grants from the CIHR (PJT-153307 and PJT-156330).

## Author Contributions

YF, CL, JS, ND, AQ performed experiments, analyzed data, and wrote the manuscript. PB, RP, MP, KC, AT, RR, MB, DN, GL, GK, JC performed experiments. JM, DD, JD, AB, and AM analyzed the data and wrote the manuscript

The authors have no conflicts of interest to declare.

---

**Extended data Table 1:**
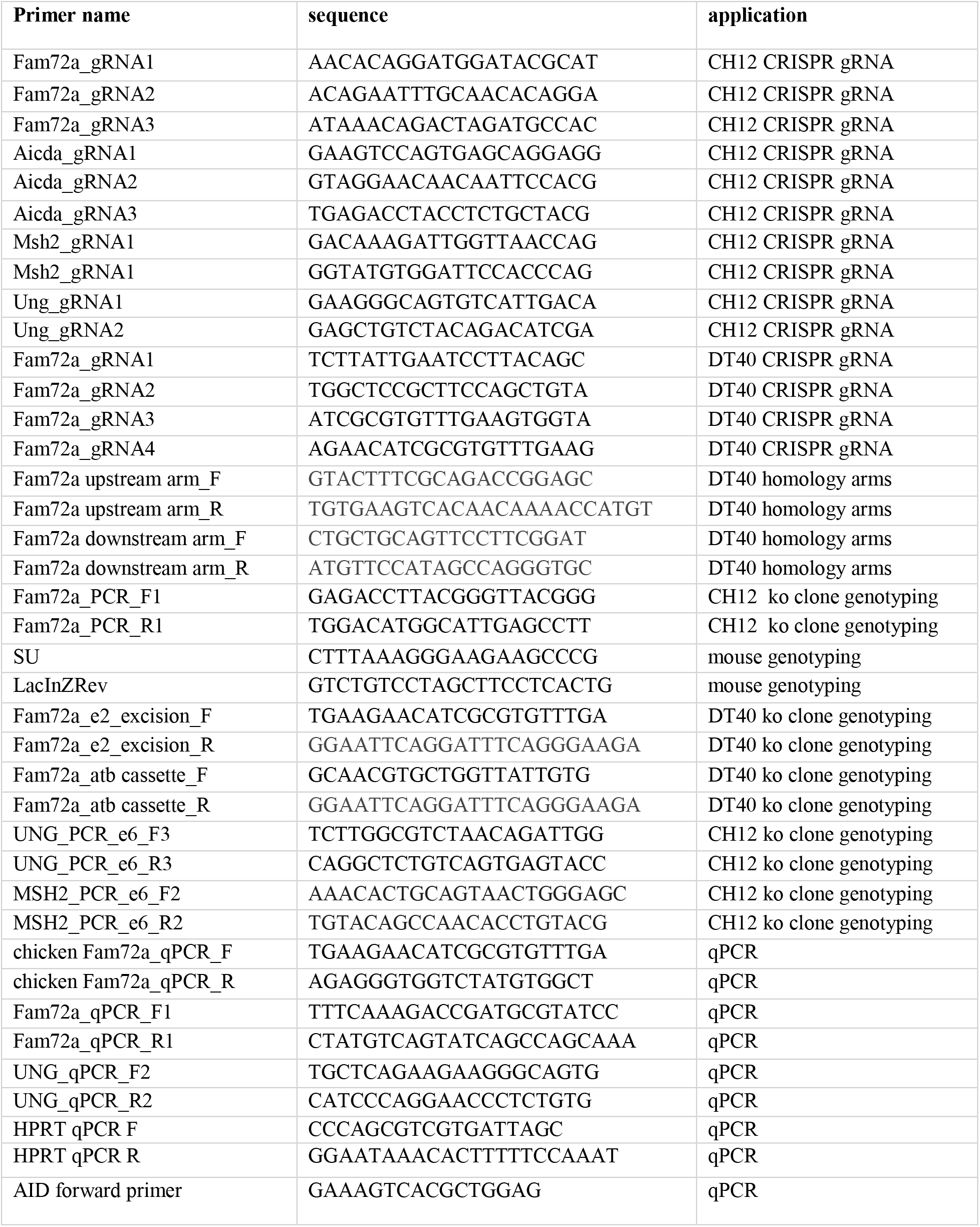
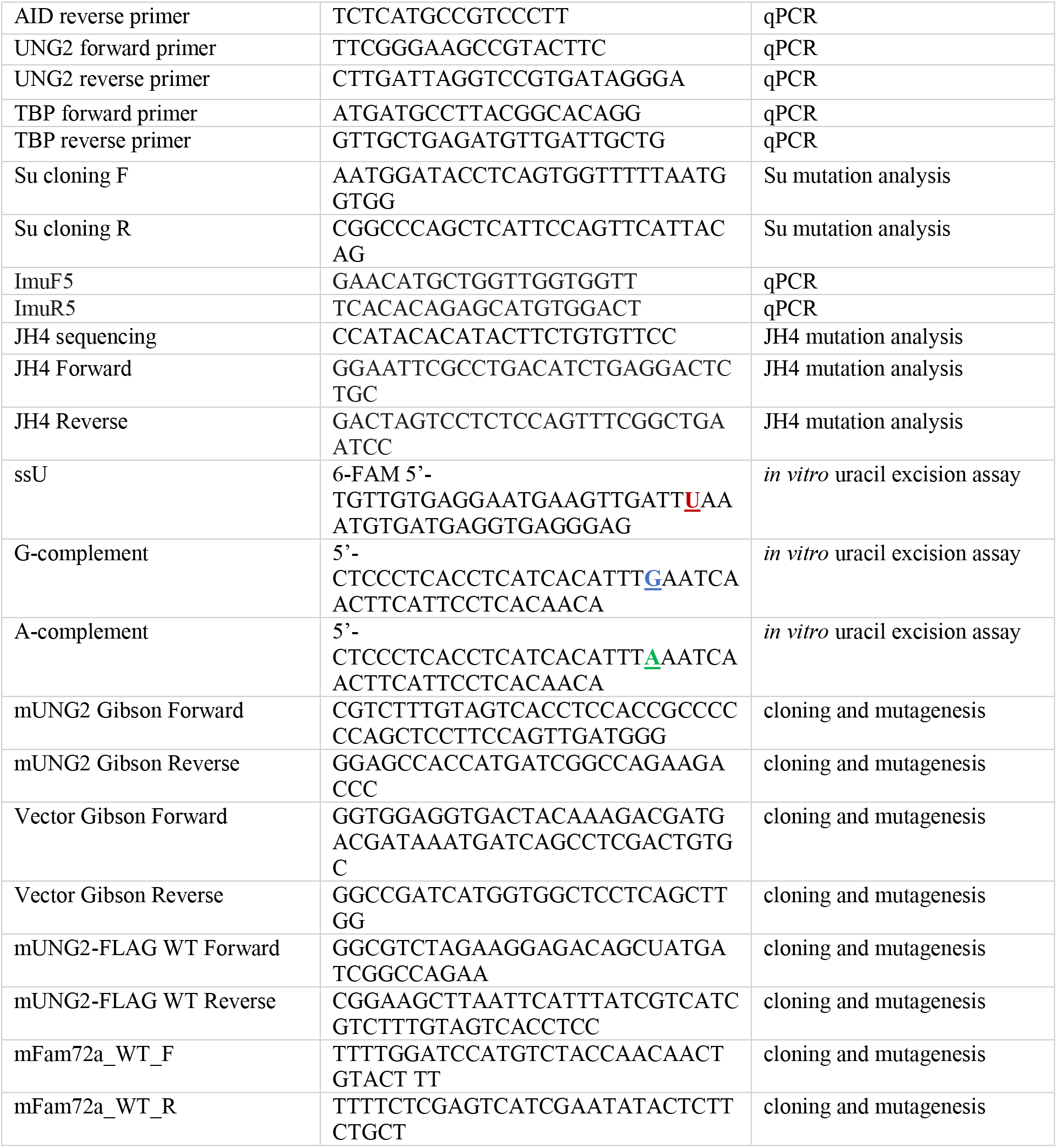
Sequences and applications for primers used in this study.

## Extended data Figure legends

**Extended data Figure 1.**
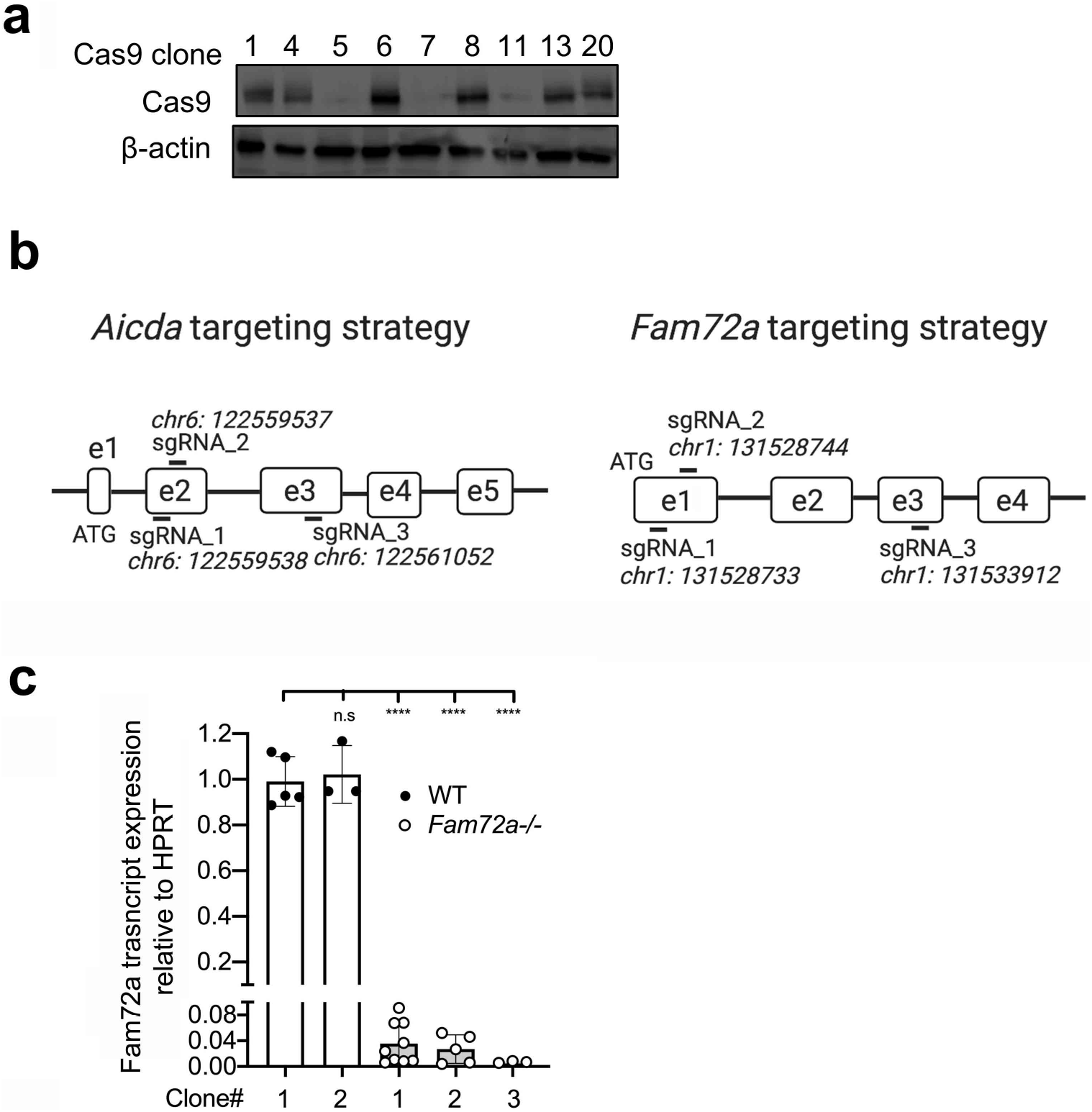
Validating the role of FAM72A during CSR in CH12 cells. (**a**) Western blot analysis to identify Cas9-expressing CH12 cell subclones after transduction with a lentiviral vector expressing Cas9. (**b**) Guide RNA (gRNA) targeting strategy against mouse *Aicda* and *Fam72a* genes to validate the role of FAM72A in CSR in bulk CH12 cells expressing Cas9. (**c**) Quantification of *Fam72a* mRNA relative to HPRT in wild-type (WT) and *Fam72a*^−/−^ CH12 clones by qPCR.

**Extended data Figure 2.**
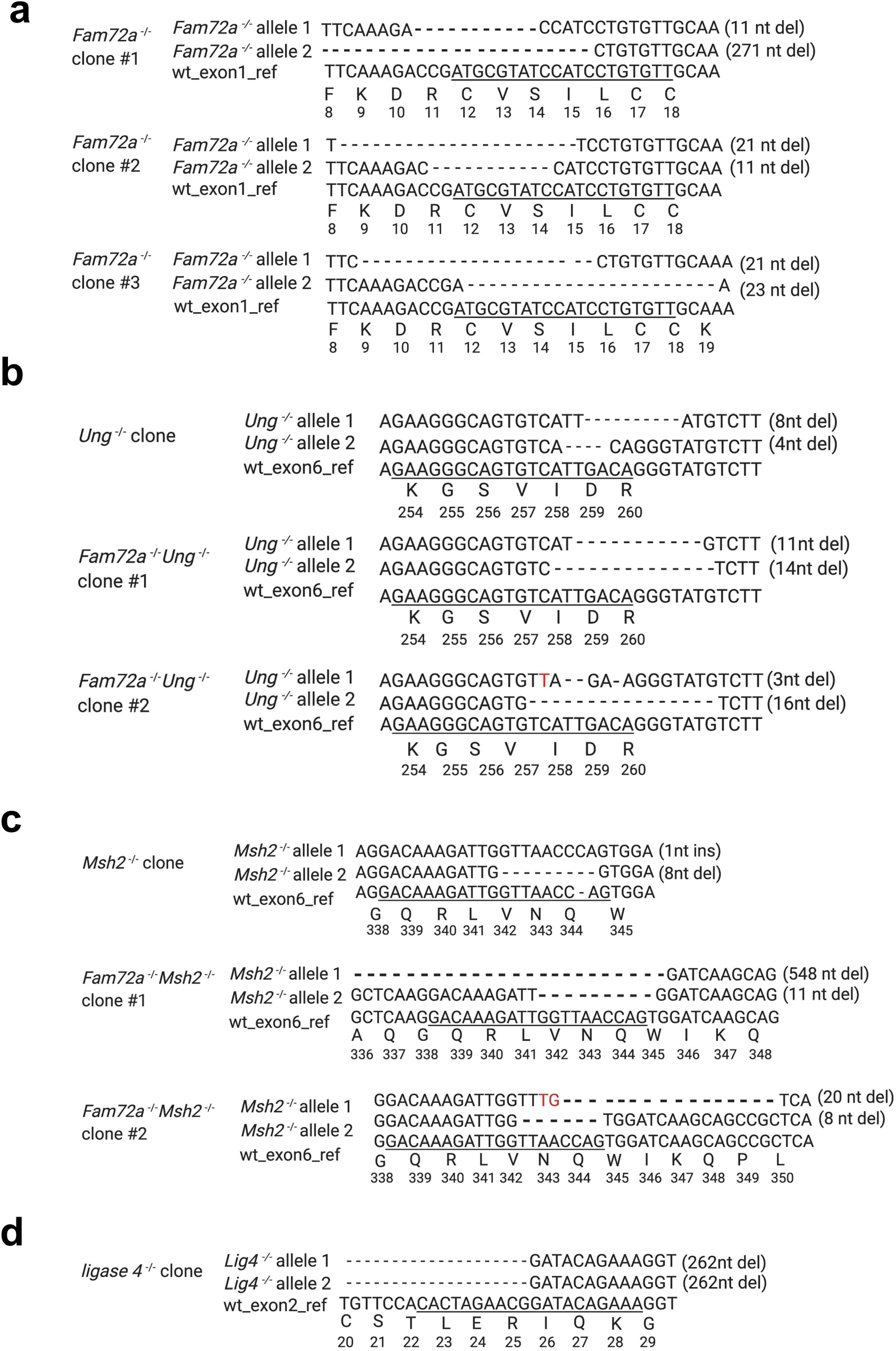
Sequenced *Fam72a, Msh2, Ung, and ligase4* alleles in *Fam72a*^−/−^,*Ung*^−/−^,*Fam72a*^−/−^*Ung*^−/−^,*Msh2^−/−^*,*Fam72a^−/−^Msh2^−/−^*, and *Ligase 4*^−/−^ CH12 clones generated using CRISPR/Cas9. Underlined sequence denotes gRNA target sequence, with the wildtype amino acid sequence indicated at the bottom.

**Extended data Figure 3.**
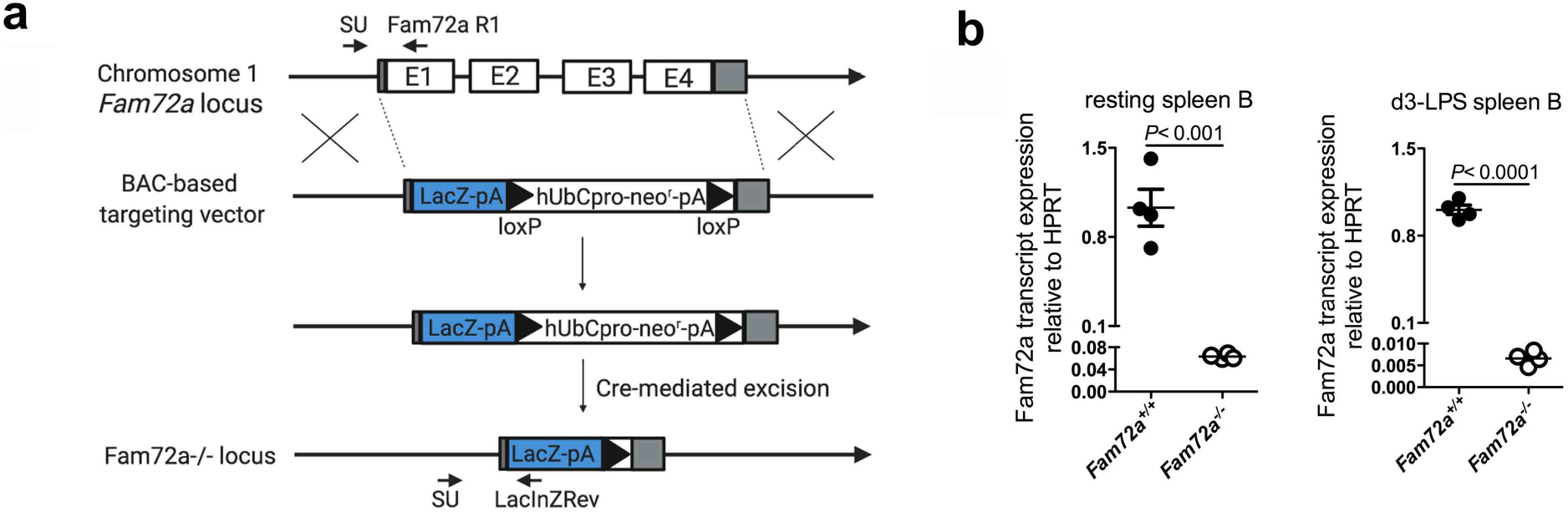
Generation of *Fam72a*^−/−^ mice. (**a**) Schematic representation of *Fam72a* gene disruption strategy. The whole coding sequence of *Fam72a* was replaced with a LacZ/neo cassette by homologous recombination using a bacterial artificial chromosome (BAC)-based targeting vector. The floxed neo cassette was removed by further breeding to a ubiquitous Cre mouse strain. Strain development was done at MMRRC (UC, Davis). PCR genotyping primer sequence can be found in **Extended data Table 1**.(**b**) qPCR analysis of *Fam72a* mRNA from resting spleen B cells or spleen B cells that were *ex vivo* stimulated with LPS for 3 days (n= 4 mice per group). Data were presented as mean ± SEM and were analyzed using two-tailed unpaired Student t test.

**Extended data Figure 4.**
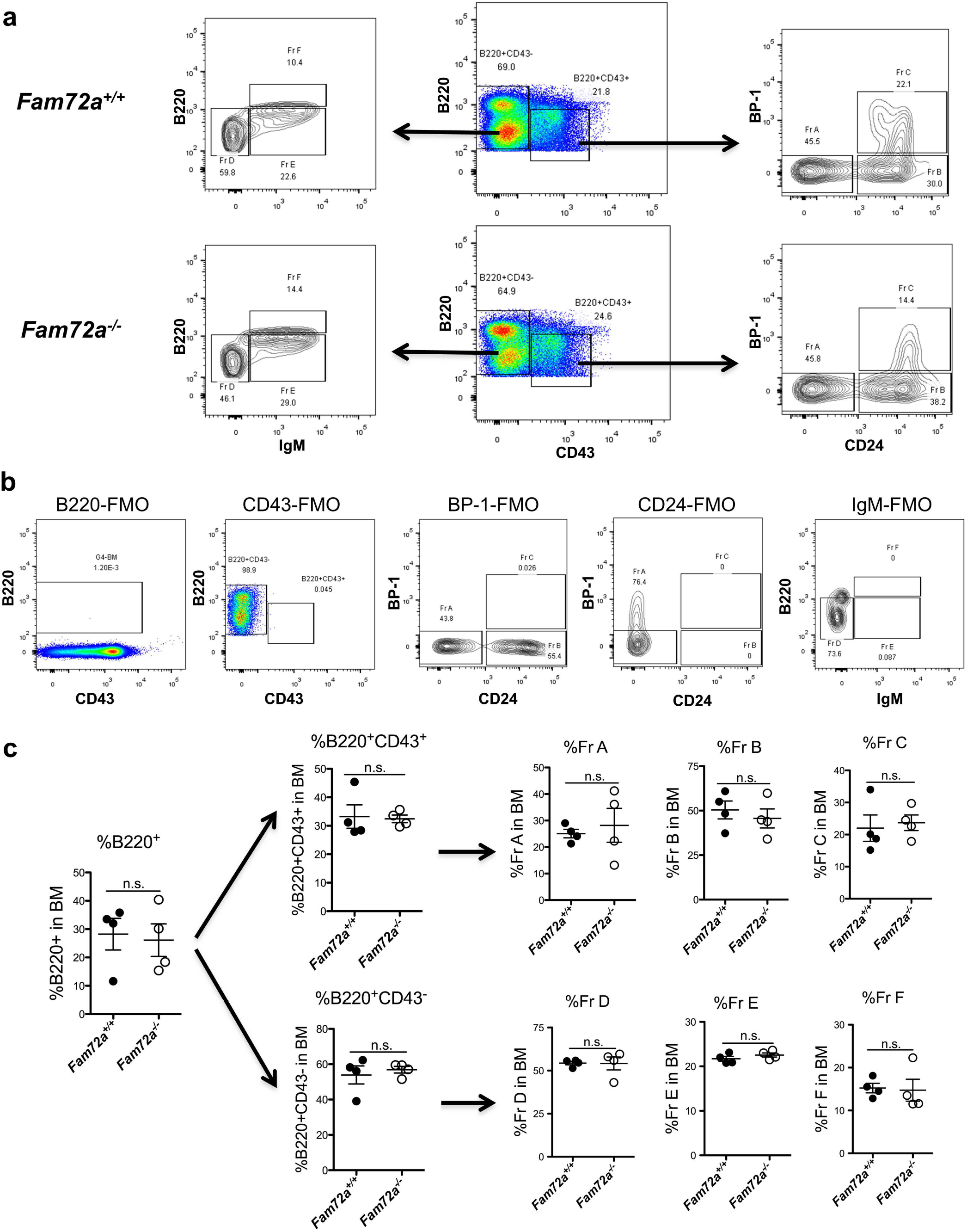
Comparable B cell profiles in the bone marrow of *Fam72a*^−/−^ and *Fam72a*^+/+^ littermate mice. (**a**) Representative FACS plots of B cells derived from the bone marrow of *Fam72a*^−/−^ or *Fam72a*^+/+^ littermate mice. (**b**) Full minus one (FMO)-derived background staining for B220, CD43, BP-1, CD24 and IgM, respectively. (**c**) The frequencies of indicated B cell fractions in the bone marrow of *Fam72a*^−/−^ or *Fam72a*^+/+^ littermates (n= 4 mice per group). Data were presented as mean ± SEM and were analyzed using two-tailed unpaired Student t test (ns: not significant). Data are representative of 2 independent experiments.

**Extended data Figure 5.**
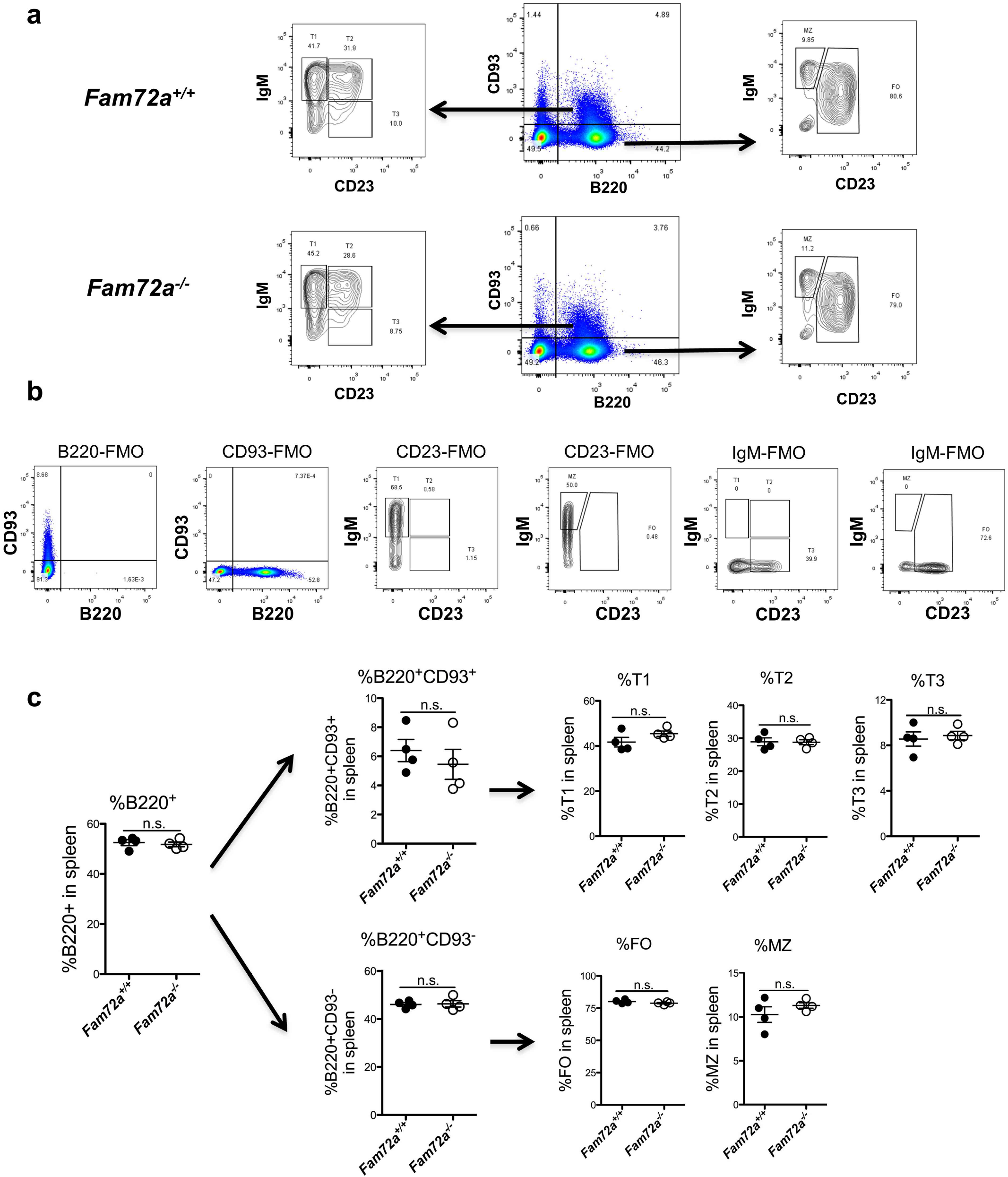
Comparable B cell profiles in the spleen of *Fam72a*^−/−^ and *Fam72a*^+/+^ littermates. (**a**) Representative FACS plots of B cells in the spleen of *Fam72a*^−/−^ or *Fam72a*^+/+^ littermate mice. (**b**) FMO-derived background staining for B220, CD93, CD23 and IgM, respectively. (**c**) The frequencies of indicated B cell subsets in the spleen of *Fam72a*^−/−^ or *Fam72a*^+/+^ littermates (n= 4 mice per group). Data were presented as mean ± SEM and were analyzed using two-tailed unpaired Student t test (ns: not significant). Data are representative of 2 independent experiments. MZ: marginal zone B cells; FO: follicular B cells.

**Extended data Figure 6.**
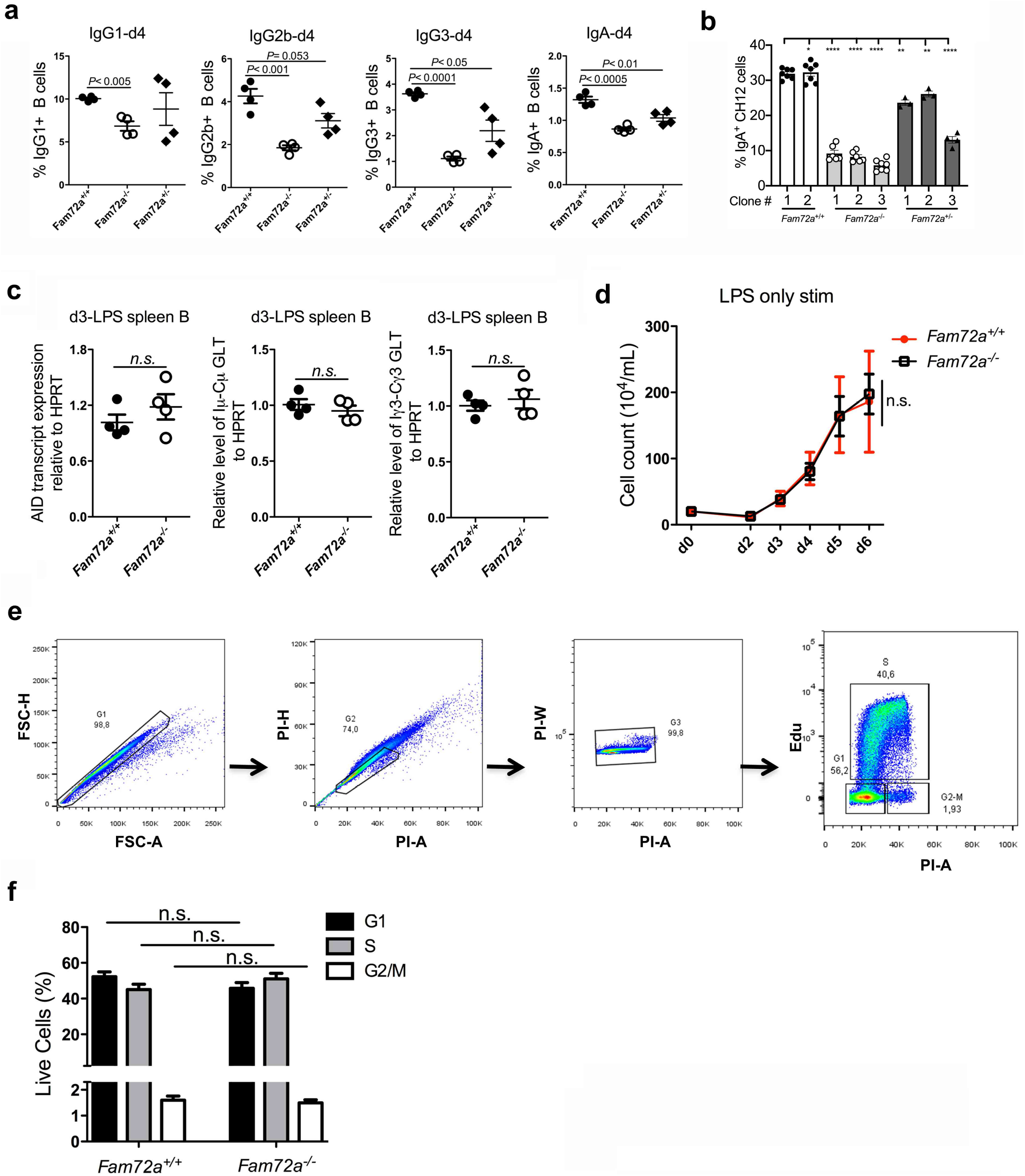
Analysis of AID and germline transcripts, cell proliferation and cell cycles of splenic B cells from *Fam72a*^−/−^ and WT littermate mice. (**a**) Analysis of *ex vivo* CSR in splenic B cells from *Fam72a*^+/+^, *Fam72a*^+/-^, and *Fam72a*^−/−^ mice. (**b**) CH12 clones of indicated genotype were treated with CIT for 2 days, and analyzed for CSR to IgA. (**c**) qPCR analysis of AID mRNA, Iμ-Cμ and Iγ3-Cγ3 germline transcripts of d3-LPS stimulated splenic B cells from *Fam72a*^−/−^ or *Fam72a*^+/+^ littermate mice. (**d**) Evaluation of cell proliferation of LPS-stimulated splenic B cells from *Fam72a*^−/−^ or *Fam72a*^+/+^ littermate mice. Data was tested by two-way ANOVA. (**e**) Gating strategy for cell cycle analysis. (**f**) The compiled cell cycle analysis of *Fam72a*^−/−^ or *Fam72a*^+/+^ splenic B cells that were stimulated by LPS for 3 days (n= 4 mice per group). Data in (a, b, and d) were presented as mean ± SEM and were analyzed using two-tailed unpaired Student t test (ns: not significant).

**Extended data Figure 7.**
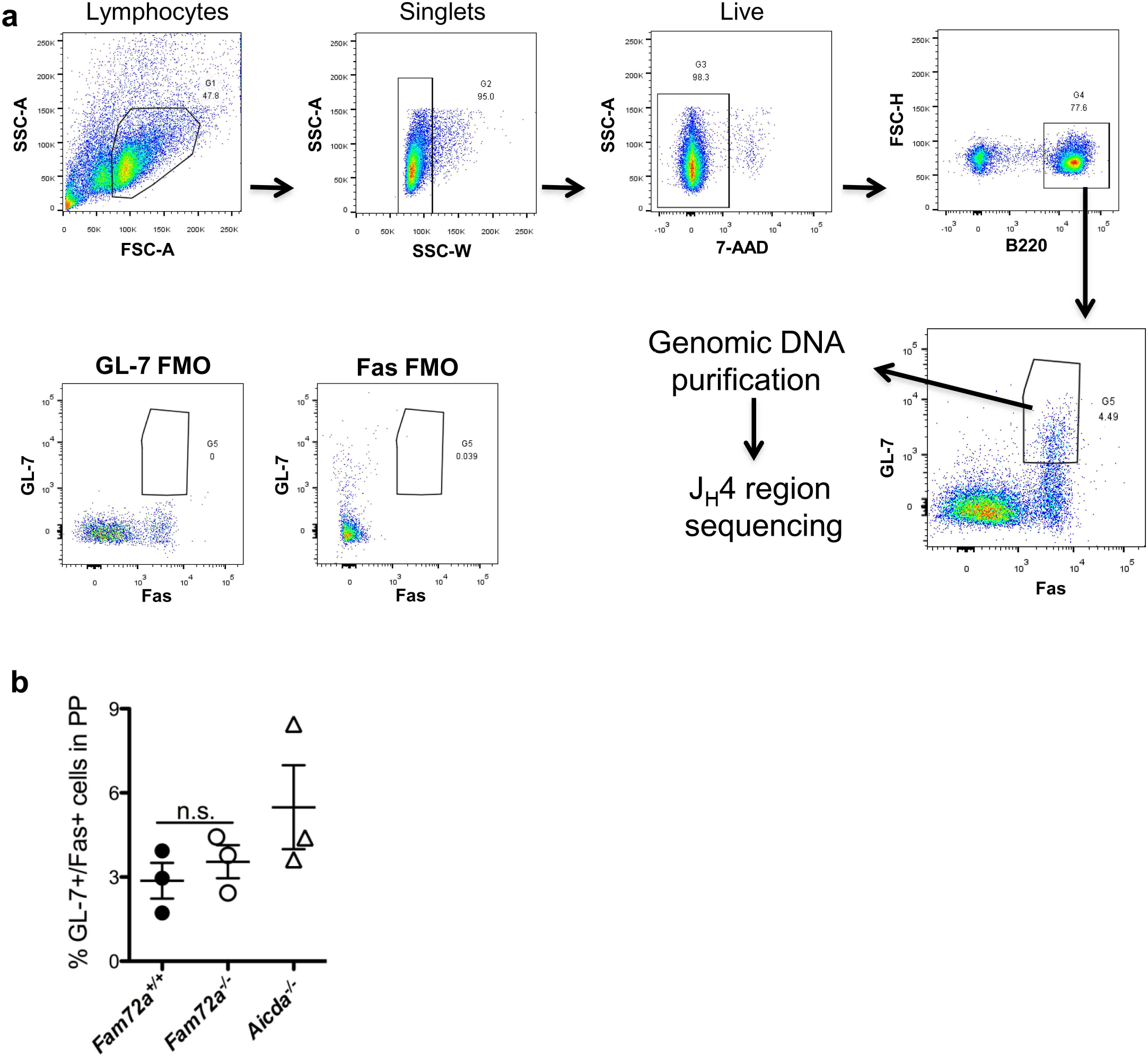
Analysis of germinal center B cells from Peyer’s patches. (**a**) Gating strategy used to sort germinal center B cells from Peyer’s patches for JH4 region sequencing. (**b**) The frequencies of germinal center B cells in Peyer’s patches from naïve *Fam72a*^+/+^, *Fam72a*^−/−^ or *Aicda*^−/−^ mice (n= 3 mice per group). Data were presented as mean ± SEM and were analyzed using two-tailed unpaired Student t test (ns: not significant).

**Extended data Figure 8.**
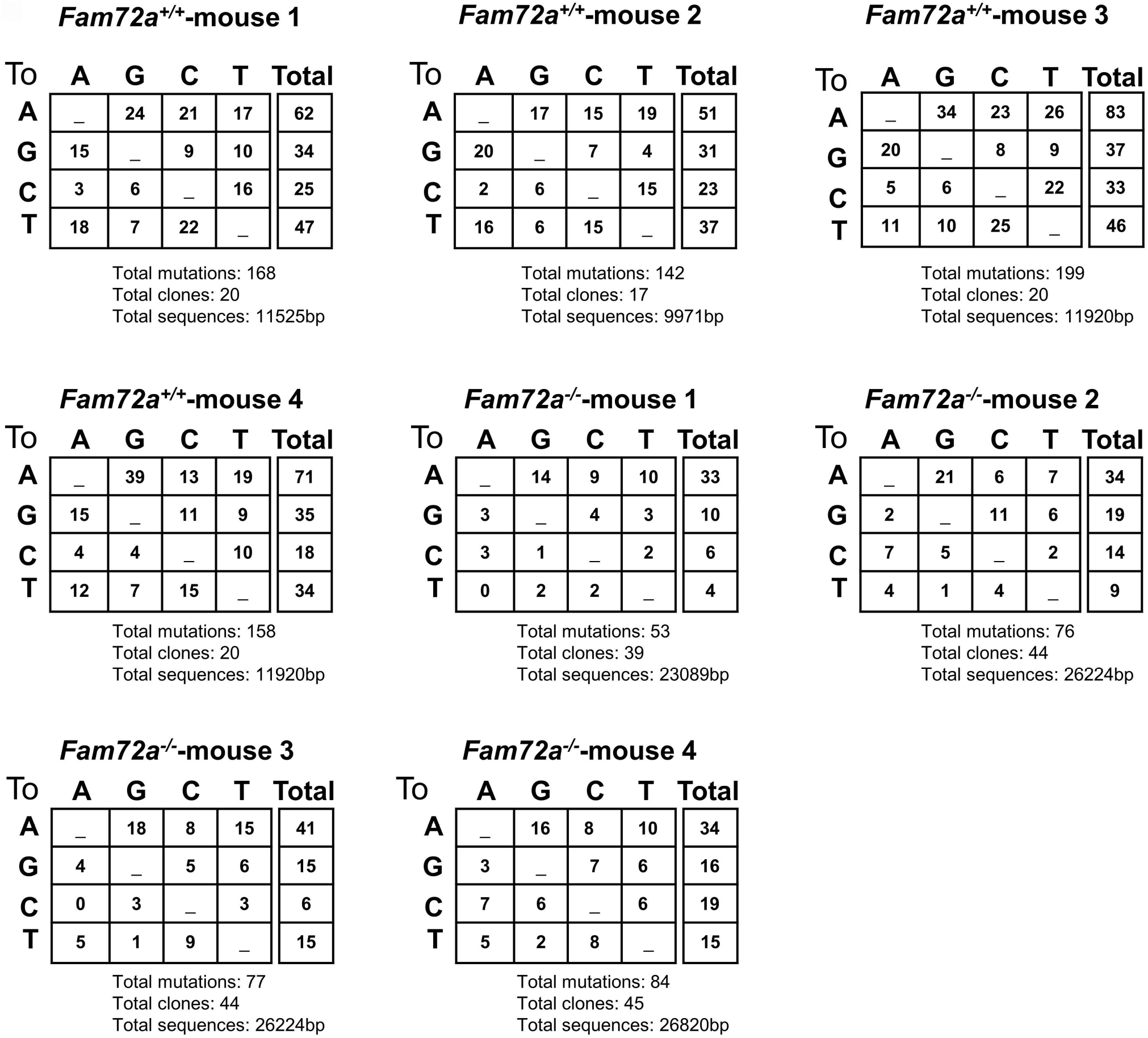
Analysis of somatic hypermutation in the JH4 region of germinal center B cells. Raw data files of JH4 region mutations of germinal center B cells for each *Fam72a*^+/+^ or *Fam72a*^−/−^ mouse.

**Extended data Figure 9.**
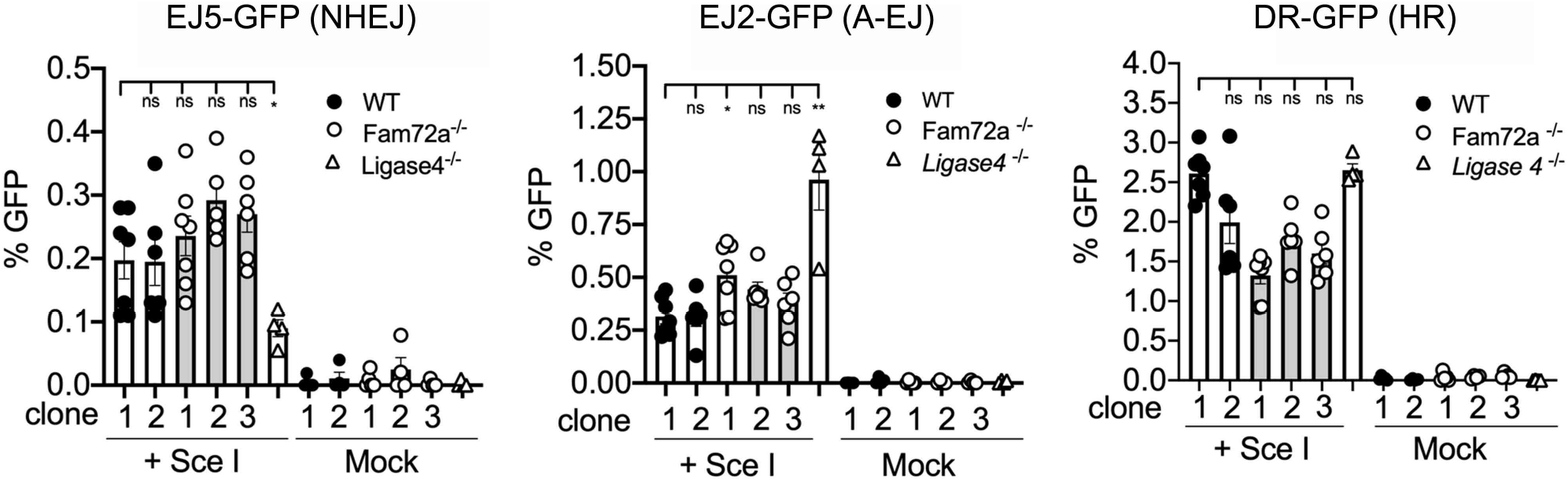
Assessing the role of FAM72A in DNA double-strand break repair pathways. WT, *Fam72a*^−/−^ and *Ligase 4*^−/−^ CH12 clones stably expressing EJ5-GPF, EJ2-GFP, and DR-GFP substrates that measure non-homologous end joining (NHEJ), alternative end joining (A-EJ), and homologous recombination (HR), respectively. Cells were mock transfected or transfected with yeast endonuclease I-SceI expressing vector, pCBA-SceI. GFP expression was monitored by flow cytometry three days post-transfection. Data are representative of three independent experiments. *, p<0.05; **, p<0.01; ***, p<0.001; ****, p<0.0001.

**Extended data Figure 10.**
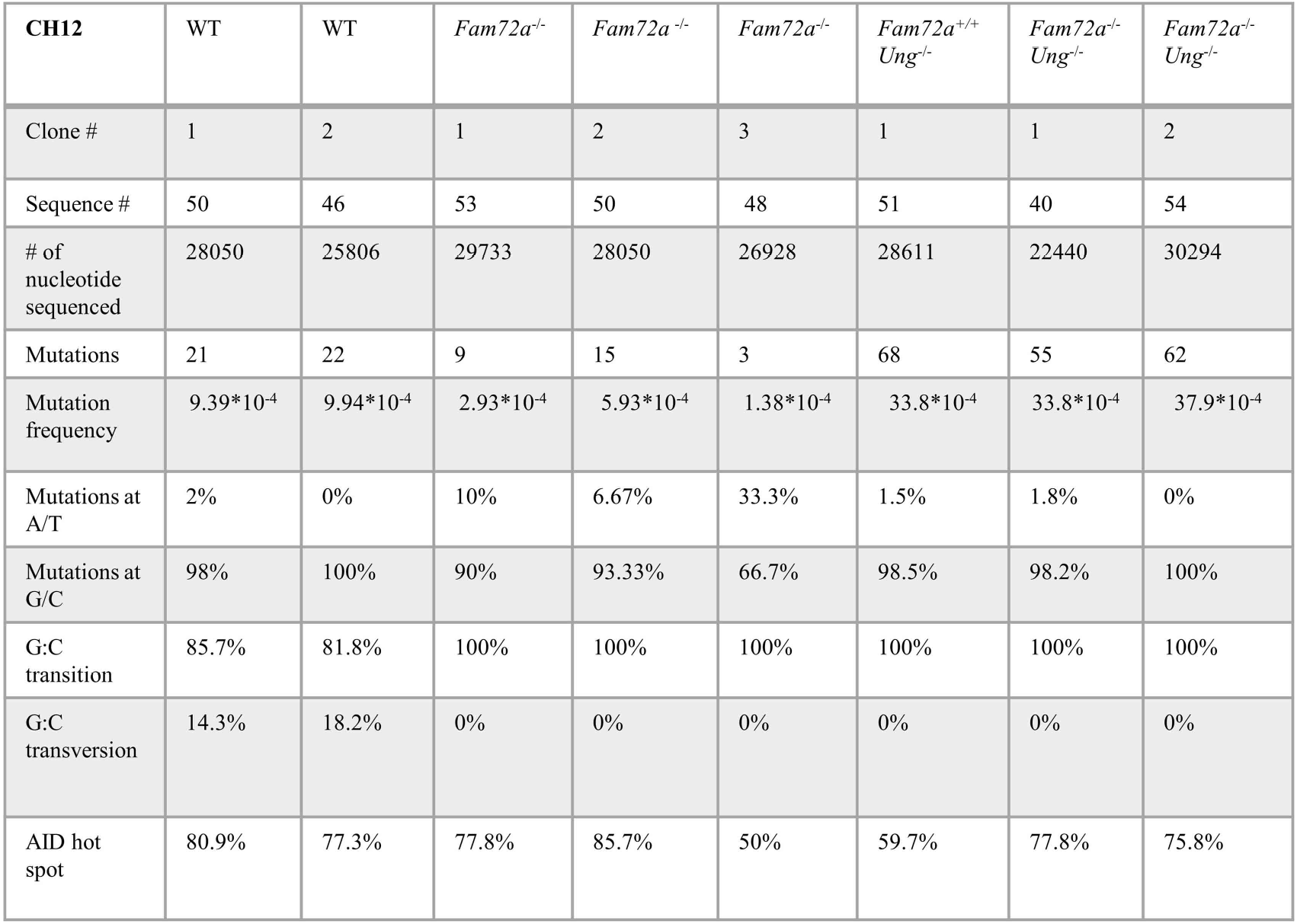
Evaluating the effect of UNG- and Msh2-deficiency on mutations in the 5’Sμ region in *Fam72a*−/− CH12 cells. Mutation characteristics at the 5’ Sμ region was analyzed in WT, *Fam72a^−/−^, Ung^−/−^*, and *Ung*^−/−^ *Fam72a*^−/−^ CH12 clones after 5 days of CIT treatment. Mutations at WRC and GYW motifs are considered AID hotspot mutation, where W=A/T, R=A/G, and Y=C/T. Mutation frequency was calculated from total mutations pooled from three independent experiment divided by total number of nucleotide sequenced. Data is summarized in **Fig. 3a**.

**Extended data Figure 11.**
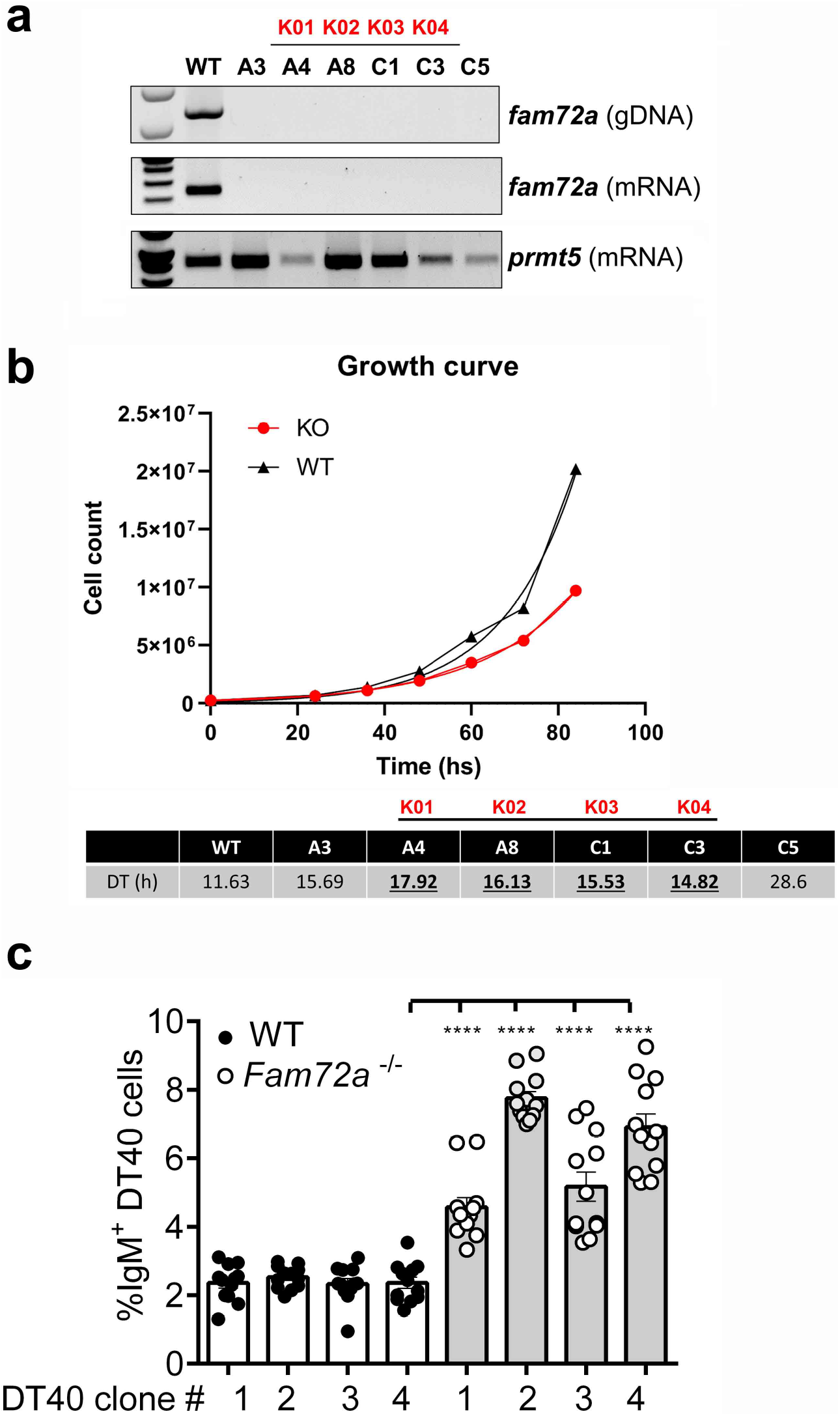
Characterization of chicken DT40 *Fam72a*^−/−^ clones. (**a**) *Fam72a*^−/−^ DT40 clones were generated using CRISPR/Cas9. Depletion of *Fam72a* was confirmed at the genomic DNA and mRNA level in 6 different DT40 clones. All the clones were mostly IgM^−^ as assessed by flow cytometry and only 4 clones (renames K01-04) were picked for fluctuation analysis. *Prmt5* was used as a control for mRNA extraction. (**b**) Growth curve analysis in WT and *Fam72a*^−/−^ DT40 cells. The average doubling time (DT) of WT and *Fam72a*^−/−^ clones in cultures for each individual clone is shown in the bottom panel and revealed reduced growth kinetics for all *Fam72a*^−/−^ clones (**c**) Fluctuation analysis for gene conversion in WT and *Fam72a*^−/−^ DT40 cells based on same number of cell divisions.

**Extended data Figure 12.**
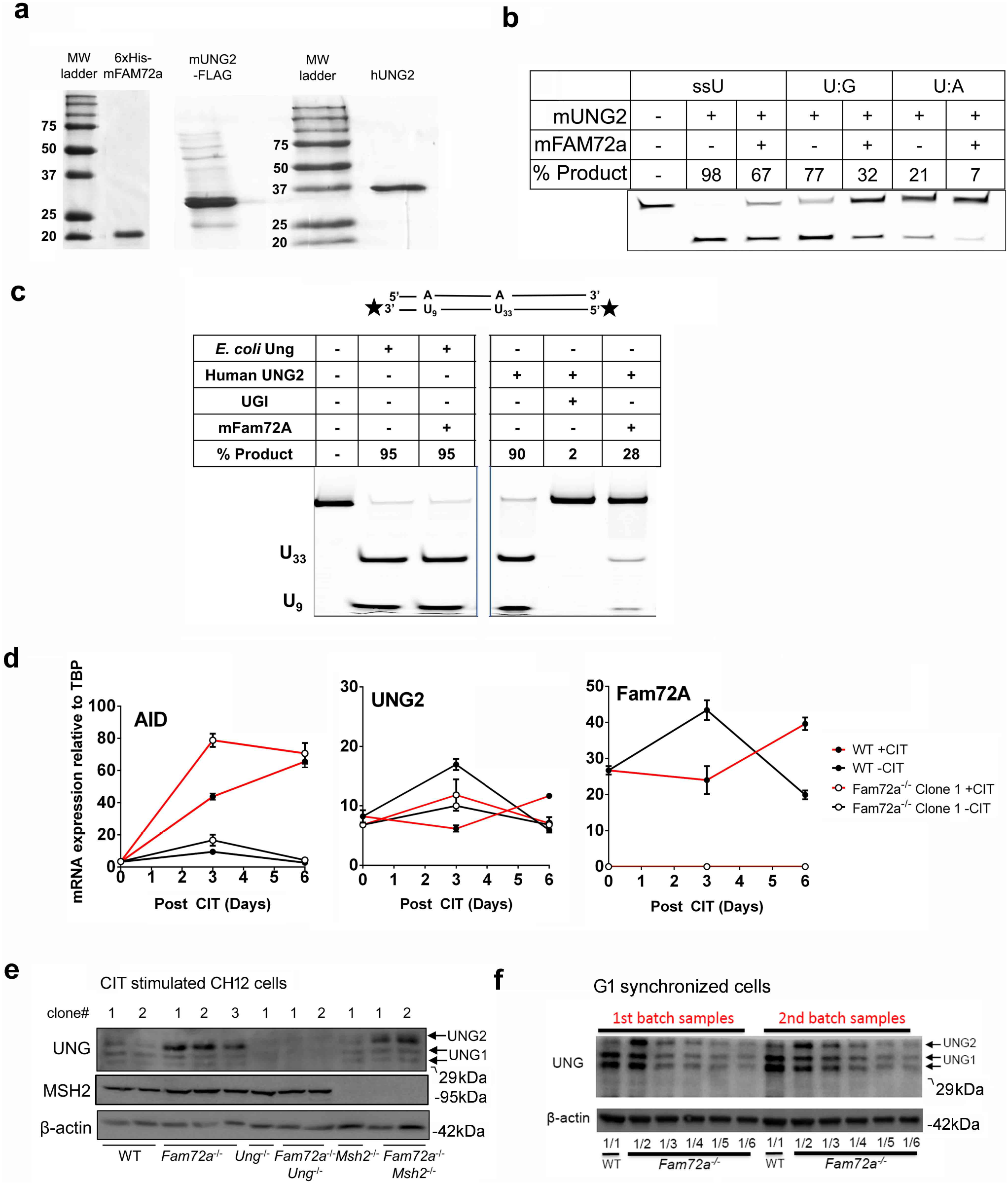
Purification and enzymatic analysis of mFAM72A on mUNG2 and hUNG2 activity, and UNG2 protein levels. (**a**) Purified mFAM72A and UNG2 proteins were electrophoresed on an SDS-PAGE gel and stained with Coomassie Brilliant Blue dye. The 6XHis-mFAM72A (20 kDa), mUNG2-polyGly-FLAG (35kDa), and hUNG2 (35kDa) are seen next to the precision plus protein standard (Bio-Rad). (**b**) Purified murine UNG2 and Fam72A proteins were pre-incubated then reacted with either a single-stranded uracil substrate (ssU) or double-stranded uracil substrates with either U:G or U:A pairs. Reaction products were separated on a denaturing polyacrylamide gel. Data is summarized in **Fig. 4c**. (**c**) Inhibition of hUNG2, but not *E. coli* UNG by mFAM72A. Purified *E. coli* UNG or hUNG2 proteins were pre-incubated with a five-fold molar excess of mFAM72A and then reacted with a double-stranded DNA substrate, with uracils at position 9 and 33 in a 55 bp DNA. Both the ends of the uracil-containing oligomer are labeled with 6-FAM (shown as stars). Consequently, two major excision products (U9 and U33) are observed. Uracil glycosylase inhibitor (Ugi) was used in one of the reactions. (**d**) AID, UNG2 and FAM72A expression in CH12 cell lines. The mRNAs for *Aicda, Ung2*, and *Fam72a* genes were quantified from WT (clone 1) and *Fam72a*^−/−^ (clone 1) cell lines that had been stimulated with the CIT cocktail (+CIT) or untreated (-CIT). Mean values and standard deviations of three independent qRTPCR reactions are shown for each treatment. The expression was normalized to the reference gene *TBP*. (**e**) Western blots for UNG, MSH2, and β-ACTIN (as control) in CIT-stimulated CH12 clones of the indicated genotype. UNG1 and UNG2 are indicated on the gel. (**f**) Lysates from G1-synchronized *Fam72a*^−/−^ CH12 cells were diluted to determine the level of increase of UNG2 protein compared to undiluted WT CH12 cells, then probed for UNG protein by western blot. These blots suggest a 3-4 fold increase in UNG2 protein in *Fam72a*^−/−^ CH12 cells compared to controls.

**Extended data Figure 13.**
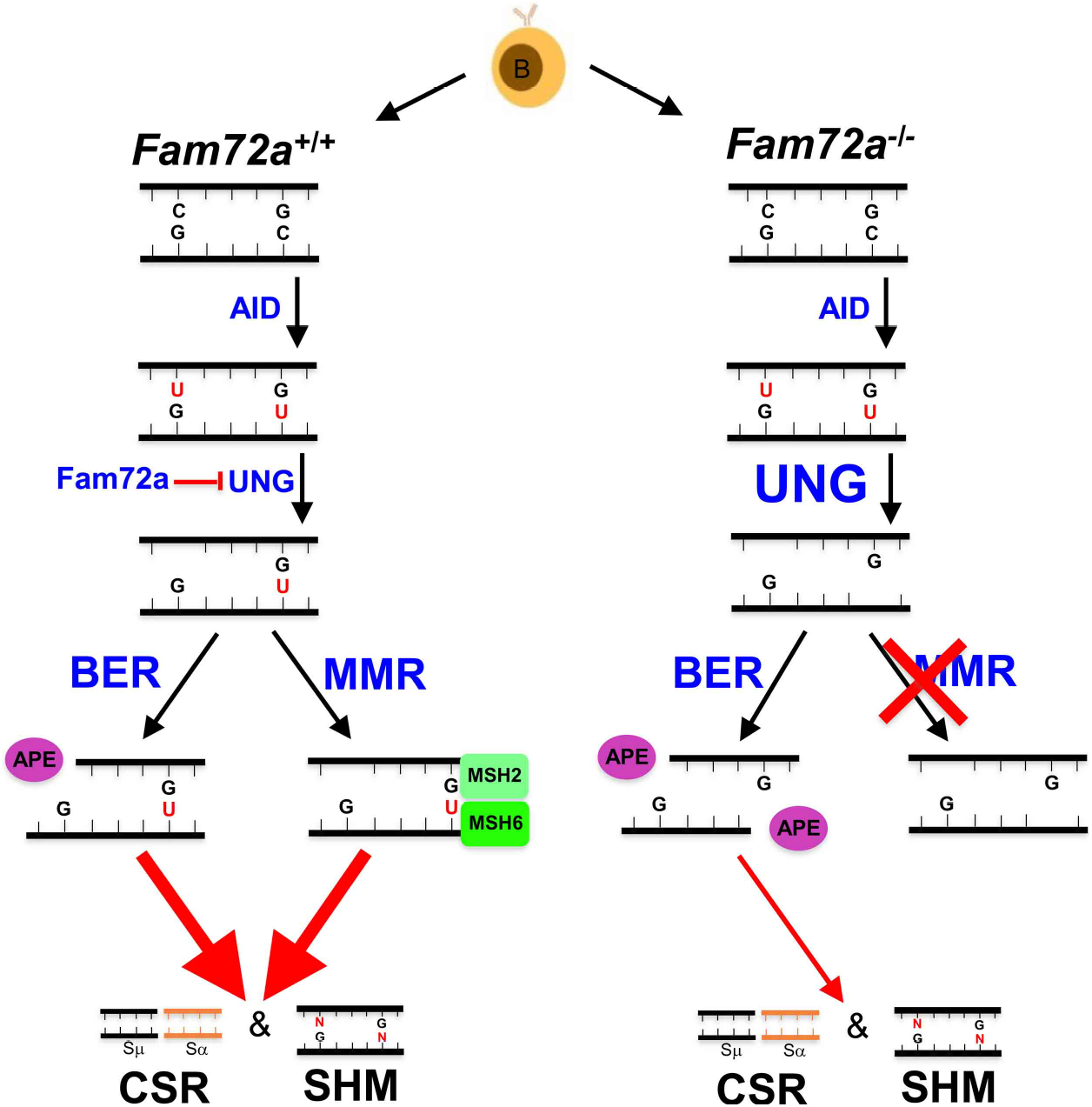
Model for the role of FAM72A in SHM and CSR. FAM72A antagonizes UNG2 in the G1 phase of the cell cycle, leading to reduced processing of the AID-induced dU in *Ig* genes, which leads to increased G:U mismatches. The increased G:U lesions can either be replicated to produce transition mutations at G:C basepairs, or can engage the mismatch repair system to enhance mutagenesis, or provide DNA lesions, in collaboration with UNG2 that are required for CSR. In the absence of FAM72A, the protein expression and enzymatic activity of UNG2 are enhanced, resulting in expedited excision of dU, which favors the faithful repair by base excision repair pathway, as evidenced by diminished mutation frequencies in both μ switch region and JH4 region. Furthermore, the accelerated removal of dU in the context of FAM72A deficiency fails to activate mismatch repair system, which is required for both CSR and SHM

## References

1 Muramatsu, M. et al. Class switch recombination and hypermutation require activation-induced cytidine deaminase (AID), a potential RNA editing enzyme. Cell 102, 553–563 (2000).

2 Feng, Y., Seija, N., Jm, D. I. N. & Martin, A. AID in Antibody Diversification: There and Back Again. Trends Immunol 41, 586–600, doi:10.1016/j.it.2020.04.009 (2020).

3 Cascalho, M., Wong, J., Steinberg, C. & Wabl, M. Mismatch repair co-opted by hypermutation. Science 279, 1207–1210, doi:10.1126/science.279.5354.1207 (1998).

4 Martin, A. & Scharff, M. D. AID and mismatch repair in antibody diversification. Nature reviews. Immunology 2, 605–614, doi:10.1038/nri858 (2002).

5 Phung, Q. H. et al. Increased hypermutation at G and C nucleotides in immunoglobulin variable genes from mice deficient in the MSH2 mismatch repair protein. The Journal of experimental medicine 187, 1745–1751, doi:10.1084/jem.187.11.1745 (1998).

6 Di Noia, J. & Neuberger, M. S. Altering the pathway of immunoglobulin hypermutation by inhibiting uracil-DNA glycosylase. Nature 419, 43–48, doi:10.1038/nature00981 (2002).

7 Wiesendanger, M., Kneitz, B., Edelmann, W. & Scharff, M. D. Somatic hypermutation in MutS homologue (MSH)3-, MSH6-, and MSH3/MSH6-deficient mice reveals a role for the MSH2-MSH6 heterodimer in modulating the base substitution pattern. The Journal of experimental medicine 191, 579–584, doi:10.1084/jem.191.3.579 (2000).

8 Rada, C., Di Noia, J. M. & Neuberger, M. S. Mismatch recognition and uracil excision provide complementary paths to both Ig switching and the A/T-focused phase of somatic mutation. Molecular cell 16, 163–171, doi:10.1016/j.molcel.2004.10.011 (2004).

9 Rada, C. et al. Immunoglobulin isotype switching is inhibited and somatic hypermutation perturbed in UNG-deficient mice. Current biology : CB 12, 1748–1755, doi:10.1016/s0960-9822(02)01215-0 (2002).

10 Yu, K. & Lieber, M. R. Current insights into the mechanism of mammalian immunoglobulin class switch recombination. Crit Rev Biochem Mol Biol 54, 333–351, doi:10.1080/10409238.2019.1659227 (2019).

11 Frieder, D., Larijani, M., Collins, C., Shulman, M. & Martin, A. The concerted action of Msh2 and UNG stimulates somatic hypermutation at A. T base pairs. Molecular and cellular biology 29, 5148–5157, doi:10.1128/MCB.00647-09 (2009).

12 Thientosapol, E. S. et al. Proximity to AGCT sequences dictates MMR-independent versus MMR-dependent mechanisms for AID-induced mutation via UNG2. Nucleic acids research 45, 3146–3157, doi:10.1093/nar/gkw1300 (2017).

13 Girelli Zubani, G. et al. Pms2 and uracil-DNA glycosylases act jointly in the mismatch repair pathway to generate Ig gene mutations at A-T base pairs. The Journal of experimental medicine 214, 1169–1180, doi:10.1084/jem.20161576 (2017).

14 Muramatsu, M. et al. Specific expression of activation-induced cytidine deaminase (AID), a novel member of the RNA-editing deaminase family in germinal center B cells. The Journal of biological chemistry 274, 18470–18476, doi:10.1074/jbc.274.26.18470 (1999).

15 Lawson, K. A. et al. Functional genomic landscape of cancer-intrinsic evasion of killing by T cells. Nature 586, 120–126, doi:10.1038/s41586-020-2746-2 (2020).

16 Guo, C. et al. Ugene, a newly identified protein that is commonly overexpressed in cancer and binds uracil DNA glycosylase. Cancer research 68, 6118–6126, doi:10.1158/0008-5472.CAN-08-1259 (2008).

17 Perez-Duran, P. et al. UNG shapes the specificity of AID-induced somatic hypermutation. The Journal of experimental medicine 209, 1379–1389, doi:10.1084/jem.20112253 (2012).

18 Di Noia, J. M., Rada, C. & Neuberger, M. S. SMUG1 is able to excise uracil from immunoglobulin genes: insight into mutation versus repair. The EMBO journal 25, 585–595, doi:10.1038/sj.emboj.7600939 (2006).

19 Manis, J. P. et al. 53BP1 links DNA damage-response pathways to immunoglobulin heavy chain class-switch recombination. Nature immunology 5, 481–487, doi:10.1038/ni1067 (2004).

20 Pan-Hammarstrom, Q. et al. Impact of DNA ligase IV on nonhomologous end joining pathways during class switch recombination in human cells. The Journal of experimental medicine 201, 189–194, doi:10.1084/jem.20040772 (2005).

21 Ward, I. M. et al. 53BP1 is required for class switch recombination. J Cell Biol 165, 459–464, doi:10.1083/jcb.200403021 (2004).

22 Yan, C. T. et al. IgH class switching and translocations use a robust non-classical end-joining pathway. Nature 449, 478–482, doi:10.1038/nature06020 (2007).

23 Arakawa, H., Saribasak, H. & Buerstedde, J. M. Activation-induced cytidine deaminase initiates immunoglobulin gene conversion and hypermutation by a common intermediate. PLoS Biol 2, E179, doi:10.1371/journal.pbio.0020179 (2004).

24 Di Noia, J. M. & Neuberger, M. S. Immunoglobulin gene conversion in chicken DT40 cells largely proceeds through an abasic site intermediate generated by excision of the uracil produced by AID-mediated deoxycytidine deamination. Eur J Immunol 34, 504–508, doi:10.1002/eji.200324631 (2004).

25 Campo, V. A. et al. MSH6- or PMS2-deficiency causes re-replication in DT40 B cells, but it has little effect on immunoglobulin gene conversion or on repair of AID-generated uracils. Nucleic acids research 41, 3032–3046, doi:10.1093/nar/gks1470 (2013).

26 Wang, Q. et al. The cell cycle restricts activation-induced cytidine deaminase activity to early G1. The Journal of experimental medicine 214, 49–58, doi:10.1084/jem.20161649 (2017).

27 Cappelli, E. et al. Rates of base excision repair are not solely dependent on levels of initiating enzymes. Carcinogenesis 22, 387–393, doi:10.1093/carcin/22.3.387 (2001).

28 Krusong, K., Carpenter, E. P., Bellamy, S. R., Savva, R. & Baldwin, G. S. A comparative study of uracil-DNA glycosylases from human and herpes simplex virus type 1. The Journal of biological chemistry 281, 4983–4992, doi:10.1074/jbc.M509137200 (2006).

29 Traut, T. W. Physiological concentrations of purines and pyrimidines. Mol Cell Biochem 140, 1–22, doi:10.1007/BF00928361 (1994).

30 Siriwardena, S. U., Perera, M. L. W., Senevirathne, V., Stewart, J. & Bhagwat, A. S. A Tumor-Promoting Phorbol Ester Causes a Large Increase in APOBEC3A Expression and a Moderate Increase in APOBEC3B Expression in a Normal Human Keratinocyte Cell Line without Increasing Genomic Uracils. Molecular and cellular biology 39, doi:10.1128/MCB.00238-18 (2019).

31 Galashevskaya, A. et al. A robust, sensitive assay for genomic uracil determination by LC/MS/MS reveals lower levels than previously reported. DNA Repair (Amst) 12, 699–706, doi:10.1016/j.dnarep.2013.05.002 (2013).

32 Zeng, X. et al. DNA polymerase eta is an A-T mutator in somatic hypermutation of immunoglobulin variable genes. Nature immunology 2, 537–541, doi:10.1038/88740 (2001).

33 Hagen, L. et al. Cell cycle-specific UNG2 phosphorylations regulate protein turnover, activity and association with RPA. The EMBO journal 27, 51–61, doi:10.1038/sj.emboj.7601958 (2008).

34 Rahane, C. S., Kutzner, A. & Heese, K. A cancer tissue-specific FAM72 expression profile defines a novel glioblastoma multiform (GBM) gene-mutation signature. J Neurooncol 141, 57–70, doi:10.1007/s11060-018-03029-3 (2019).

35 Kutzner, A., Pramanik, S., Kim, P. S. & Heese, K. All-or-(N)One - an epistemological characterization of the human tumorigenic neuronal paralogous FAM72 gene loci. Genomics 106, 278–285, doi:10.1016/j.ygeno.2015.07.003 (2015).

36 Shalhout, S. et al. Genomic uracil homeostasis during normal B cell maturation and loss of this balance during B cell cancer development. Molecular and cellular biology 34, 4019–4032, doi:10.1128/MCB.00589-14 (2014).

## Methods References

1 Ramachandran, S. et al. The SAGA Deubiquitination Module Promotes DNA Repair and Class Switch Recombination through ATM and DNAPK-Mediated gammaH2AX Formation. Cell reports 15, 1554–1565, doi:10.1016/j.celrep.2016.04.041 (2016).

2 Lawson, K. A. et al. Functional genomic landscape of cancer-intrinsic evasion of killing by T cells. Nature 586, 120–126, doi:10.1038/s41586-020-2746-2 (2020).

3 Aregger, M., Chandrashekhar, M., Tong, A. H. Y., Chan, K. & Moffat, J. Pooled Lentiviral CRISPR-Cas9 Screens for Functional Genomics in Mammalian Cells. Methods Mol Biol 1869, 169–188, doi:10.1007/978-1-4939-8805-1_15 (2019).

4 Sarno, A. et al. Uracil-DNA glycosylase UNG1 isoform variant supports class switch recombination and repairs nuclear genomic uracil. Nucleic acids research 47, 4569–4585, doi:10.1093/nar/gkz145 (2019).

5 Campo, V. A. et al. MSH6- or PMS2-deficiency causes re-replication in DT40 B cells, but it has little effect on immunoglobulin gene conversion or on repair of AID-generated uracils. Nucleic acids research 41, 3032–3046, doi:10.1093/nar/gks1470 (2013).

6 Li, C. et al. The H2B deubiquitinase Usp22 promotes antibody class switch recombination by facilitating non-homologous end joining. Nat Commun 9, 1006, doi:10.1038/s41467-018-03455-x (2018).

7 Li, C. et al. Early-life programming of mesenteric lymph node stromal cell identity by the lymphotoxin pathway regulates adult mucosal immunity. Sci Immunol 4, doi:10.1126/sciimmunol.aax1027 (2019).

8 Martin, A. et al. Msh2 ATPase activity is essential for somatic hypermutation at a-T basepairs and for efficient class switch recombination. The Journal of experimental medicine 198, 1171–1178, doi:10.1084/jem.20030880 (2003).

9 Siriwardena, S. U., Perera, M. L. W., Senevirathne, V., Stewart, J. & Bhagwat, A. S. A Tumor-Promoting Phorbol Ester Causes a Large Increase in APOBEC3A Expression and a Moderate Increase in APOBEC3B Expression in a Normal Human Keratinocyte Cell Line without Increasing Genomic Uracils. Molecular and cellular biology 39, doi:10.1128/MCB.00238-18 (2019).

10 So, C. C., Ramachandran, S. & Martin, A. E3 Ubiquitin Ligases RNF20 and RNF40 Are Required for Double-Stranded Break (DSB) Repair: Evidence for Monoubiquitination of Histone H2B Lysine 120 as a Novel Axis of DSB Signaling and Repair. Molecular and cellular biology 39, doi:10.1128/MCB.00488-18 (2019).

